# A standardized atlas of human bronchoalveolar lavage cells built using scalable ensemble annotation and cross-study robust markers

**DOI:** 10.64898/2026.01.08.698293

**Authors:** Yushan Hu, Ziying Liu, Kailun Bai, Belaid Moa, Janice M. Leung, Firoozeh V. Gerayeli, Don D. Sin, Xiaojian Shao, Xuekui Zhang

## Abstract

**Background:** Bronchoalveolar lavage (BAL) single-cell RNA sequencing (scRNA-seq) offers rich insights into pulmonary immune dynamics, yet consistent cell-type annotation remains elusive. Existing methods often rely on a single reference, risking inconsistency and domain shift across datasets. A BAL-specific, high-resolution annotation framework is critically needed.

**Methods:** We developed BAL-EA (BAL Ensemble Annotation), a BAL-centric automated annotation framework that integrates robust, cross-study marker discovery with ensemble machine learning. BAL-EA harmonizes BAL cell identities into a three-tier taxonomy (11 major lineages, 13 refined classes, 21 fine-grained subtypes) compatible with the Human Lung Cell Atlas (HLCA) while capturing lavage-enriched biology. Marker catalogues were derived via reproducibility-guided differential expression across at least 10 independent sub-studies, ensuring resilience to dataset-specific bias. Comparative benchmarking was performed against six leading annotation tools using independent BAL datasets.

**Results:** We assembled the largest BAL scRNA-seq atlas to date, integrating more than 347,333 lung cells from HLCA, multiple public BAL datasets, and the largest inhouse BAL cohort ever reported (241,924 cells from 30 individuals). BAL-EA outperformed existing annotation tools, achieving balanced macro-F1 scores over 0.95 for key lineages such as alveolar macrophages (AM), non-alveolar macrophages, and epithelial cells. Application to Chronic Obstructive Pulmonary Disease (COPD) BAL samples revealed reproducible disease-associated shifts, including increased neutrophils and CCL2-positive macrophages alongside reduced AM in COPD patients, findings validated in independent COVID-19 BAL datasets. The released atlas includes harmonized multi-resolution annotations, robust marker panels, pretrained models.

**Conclusions:** This work contributes the most comprehensive BAL scRNA-seq atlas, introduces a novel BAL-specific annotation framework (BAL-EA), standardizes BAL taxonomy at three resolutions, and provides rigorously validated marker gene resources. Together, these advances deliver a powerful reference for reproducible BAL scRNA-seq analysis and lay the foundation for clinical and translational applications in respiratory disease research.

## 1 Introduction

Single-cell RNA sequencing (scRNA-seq) of Bronchoalveolar lavage fluid (BALF) obtained via minimally invasive bronchoscopy has emerged as a powerful approach for profiling immune and epithelial cell states directly from human airspaces [1, 2]. Because BAL sampling enriches for airway/alveolar immune populations and airway-lining epithelial cells, it provides a complementary window to tissue biopsy for investigating pulmonary biology across health and disease, including Chronic Obstructive Pulmonary Disease (COPD) [3].

Recent BAL and airway-focused scRNA-seq studies have reported widely varying cell type resolutions and nomenclatures, reflecting both genuine biological heterogeneity and methodological variability. Across studies spanning healthy donors and COVID-19 patients, reported annotations range from 6 to 28 populations, with the largest divergences in myeloid and lymphoid subdivisions and their labels [4–10]. For example, Liao *et al.* identified nine main cell types in COVID-19 patients and healthy controls, including alveolar and monocyte-derived macrophages, multiple T cell subsets, and epithelial cells [4]. Wauters *et al.* expanded this to 18 populations, resolving finer subtypes of T cells, macrophages, and neutrophils [5], while Grant *et al.* described 27 immune and epithelial subtypes in severe SARS-CoV-2 pneumonia [6]. In healthy BAL, macrophages, monocytes, and T cells consistently dominate, but their subdivisions differ markedly, such as tissue-resident versus monocyte-derived macrophages or proliferative, transitional, and chemokine-enriched states, complicating cross-study comparisons and blunting the interpretability of disease-associated shifts [9, 10]. In addition, increasing the resolution of cell-type annotation through deep clustering of major immune populations can help resolve this complexity by revealing disease-relevant substructures that remain hidden at coarse annotation levels. For example, Hu *et al.* [11] demonstrated that fine-grained macrophage and monocyte stratification in diseased lungs uncovered previously unrecognized inflammatory states and highlighted potential therapeutic targets in a cell-type-specific manner.

Standardized references can further mitigate these issues by harmonizing labels, features, and annotation resolution. The recently developed Human Lung Cell Atlas (HLCA) integrates 49 tissue-based scRNA-seq studies and provides consensus annotations across major lung cell types [12]. However, the HLCA is tissue/biopsy-centric and underrepresents lavage-enriched lineages and activation states; naively projecting BAL data to tissue-derived references can yield implausible assignments or obscure BAL-dominant programs [12]. A BAL-centric atlas that remains compatible with HLCA while capturing lavage-specific biology is therefore urgently needed.

Building such a BAL atlas requires consistent, scalable, and reproducible cell-type annotation across diverse studies and cohorts. Manual annotation often guided by study-specific marker genes, becomes increasingly impractical as dataset size grows and resolution deepens, introducing variability that undermines cross-study integration. Automated annotation strategies offer a powerful alternative by enabling rapid, standardized cell type assignment, which is essential for harmonizing datasets into a coherent atlas framework. Several machine-learning-based annotation tools have been proposed, such as SingleR [13], scmap [14], CaSTLe [15], CHETAH [16], and SCINA [17]. While these general-purpose tools have demonstrated success in annotating cell types across various tissues, none of them are tailored to the BAL scRNA-seq data. More importantly, most of these annotation tools depend on predefined marker genes for cell type classification. However, robust and well-defined marker genes for diverse cell types in BAL remain largely underexplored. Furthermore, head-to-head comparisons show sensitivity to the choice of a single training reference and to cross-study technical/biological shifts (“domain shift”) [18]. In addition, leveraging existing comprehensive cell atlases, such as the HLCA dataset, would provide great potential to enhance cell type annotation.

Here we present BAL-EA, a BAL-centric framework that prioritizes robustness and standardization while remaining ontology-compatible. First, we integrate multiple BAL scRNA-seq datasets, including the largest in-house cohort to date, to define a multi-resolution BAL taxonomy that captures lavage-enriched diversity yet maps cleanly to organ-wide lineages anchored by HLCA [12]. Second, to mitigate domain shift, we derive cross-study reproducible marker catalogues by performing within-study differential analyses and selecting consensus markers, with adaptive handling of rarer populations. Third, we implement a study-wise ensemble strategy that trains per-study base classifiers on robust-marker features and aggregates them to improve stability. Benchmarking demonstrates that BAL-EA achieves high accuracy and consistency in BAL cell type assignment across diverse datasets. Finally, we release a standardized BAL atlas including marker lists, the BAL-EA framework, and curated annotations, supporting reproducible analyses and enabling direct comparisons across studies [4–6, 8–10].

## 2 Results

### 2.1 A BAL-centric three-level taxonomy and cross-study marker catalogues

We developed a standardized BAL scRNA-seq annotation framework with a three-level taxonomy (11/13/21 classes) that aligned to the HLCA [12] yet tailored to BAL data (Fig. 1). We also derived cross-study marker catalogues that are reproducible across independent cohorts. Using healthy BAL cohorts and the HLCA core sub-studies (a total of 378,113 cells), we harmonized cell type labels into Level 1 (11 major lineages), Level 2 (13 classes splitting the mononuclear phagocyte compartment), and Level 3 (21 fine subtypes capturing macrophage/monocyte programs and airway/alveolar epithelium). This hierarchy preserves the HLCA comparability while reflecting BAL-enriched biology.

**Fig. 1.**
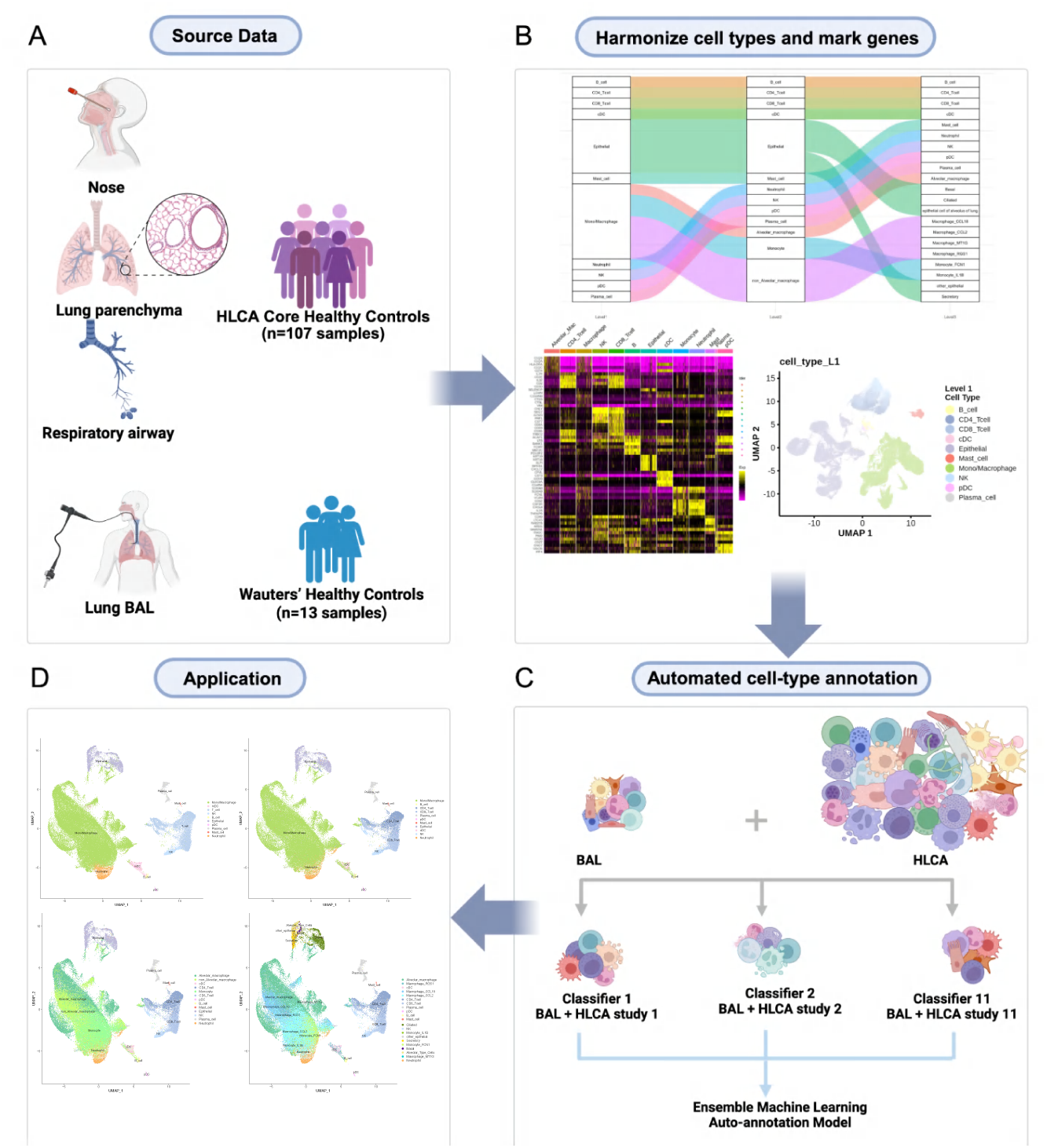
Workflow diagram of the BAL-EA. (A) Data source. BAL scRNA-seq data (n=30,780 high-quality cells from 13 healthy control subjects) and the HLCA core dataset (n=347,333 cells from 11 studies). These scRNA-seq data were integrated, quality controlled, and processed for BAL-EA training. (B) Cell type harmonization and robust marker gene identification. Common cell types were identified at various resolutions. Meanwhile, cell-type-specific genes were selected by requiring consistency across 10 out of the 11 HLCA studies. Hierarchical annotation tree (Level 1: major lineages; Level 2: broad subtypes; Level 3: fine subtypes) were generated. (C) Automated cell type annotation. Data integration through ensemble learning. (D) A comprehensive cell atlas. Multiple BAL scRNA-seq datasets were integrated together with the automated cell type annotated labels to generate a comprehensive BAL cell atlas.

At Level 1 we resolved B cells, plasma cells, CD4^+^ and CD8^+^ T cells, natural killer (NK) cells, epithelial cells, mono/macrophages, neutrophils, mast cells, conventional dendritic cells (cDCs), and plasmacytoid dendritic cells (pDCs). Level 2 refines the dominant mononuclear phagocyte compartment into AMs, non-alveolar macrophages, and monocytes while retaining other lineages (13 classes total). Level 3 adds macrophage programs (CCL18^+^, CCL2^+^, RGS1^+^, MT1G^+^), monocyte subsets (FCN1^+^, IL1B^+^, HSPA6^+^), and epithelial subtypes (basal, ciliated, secretory, alveolar), yielding 21 classes (Fig. 2A). The nomenclature for macrophage subtypes follows established conventions from prior BAL literatures on macrophage heterogeneity [4, 5, 10, 19]. Analysis of cell type composition after harmonization across datasets revealed striking variability among studies (Fig. 2B). The proportion of specific BAL cell types varied not only between independent cohorts but also within datasets derived from different disease contexts or control groups. This observation highlights the importance of harmonized reference definitions when comparing or integrating results from multiple studies.

**Fig. 2.**
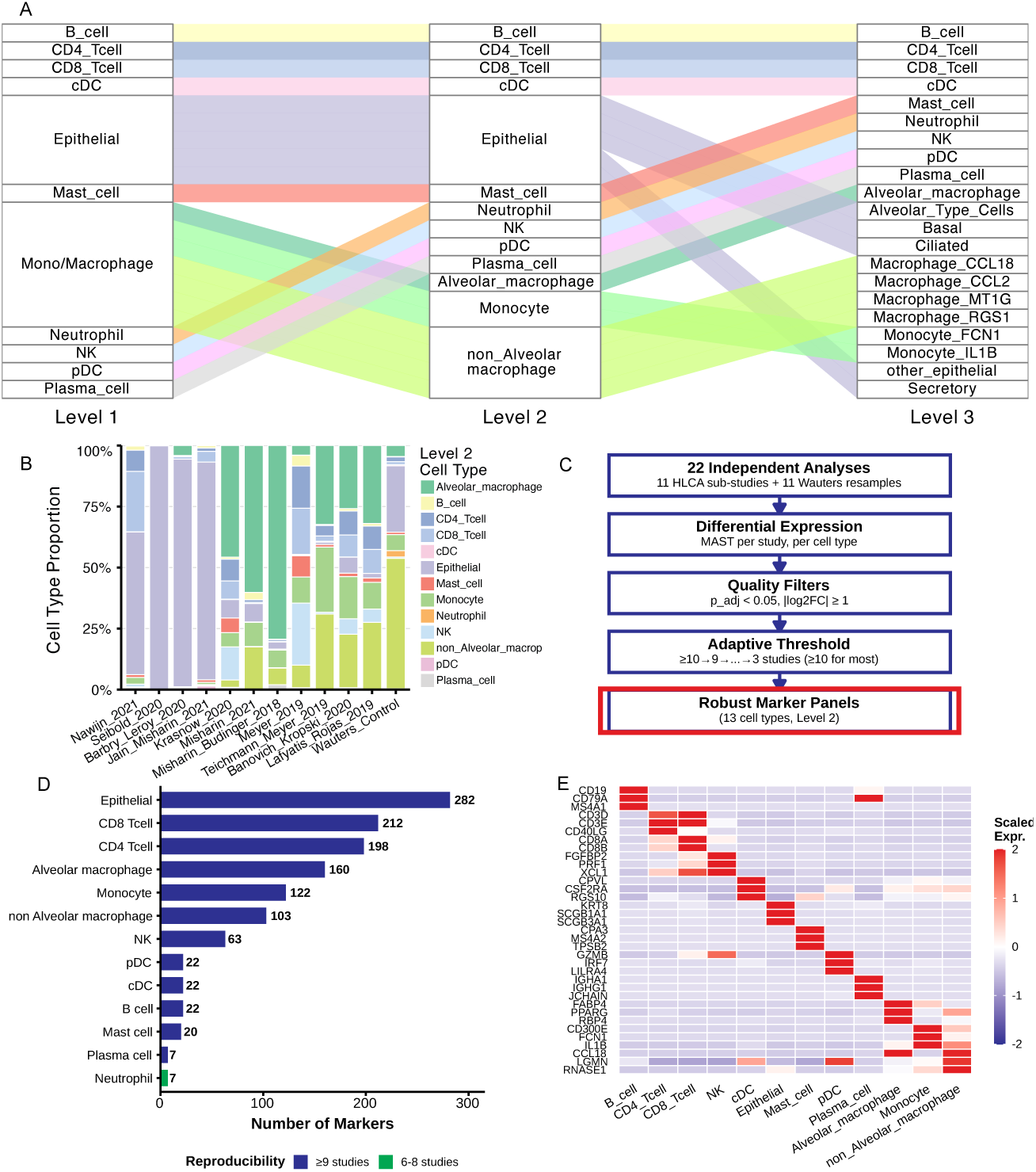
Hierarchical cell type harmonization and robust marker identification in BAL scRNA-seq Data. (A) Hierarchical cell-type structure. Sankey diagram showing relationships between Level 1 (broad lineages), Level 2 (major cell types), and Level 3 (refined subtypes). Colors indicate Level 2 cell types. (B) Cell type composition across studies. Stacked bar plot of Level 2 cell type proportions in 11 HLCA v1.0 core substudies and the Wauters *et al.* control cohort. The HLCA studies are ordered by compositional similarity, with Wauters *et al.* displayed at the end. (C) Robust marker identification pipeline. Workflow showing the multi-step approach: 22 independent DE analyses (11 HLCA substudies + 11 Wauters bootstrap resamples) → quality filters (adjusted p *<* 0.05, *|log*_2_*FC| ≥* 1) → adaptive reproducibility thresholds (*≥*10→9→…→3 studies) → consensus marker panels. (D) Marker gene expression heatmap. Scaled expression of top robust markers (up to 5 per cell type) across 13 Level 2 cell types. Red: high expression; blue: low expression. (E) Robust markers per cell type. Number of markers identified for each Level 2 cell type. Bar colors indicate reproducibility: dark blue (*≥*9 studies), green (6-8 studies). Adaptive thresholding ensured usable marker panels for both abundant and rare populations.

To obtain domain-shift-resistant features, we identified markers via within-study differential expression (DE) analysis in each HLCA sub-study and in resampled healthy Wauters *et al.* BAL controls [5], then retained genes replicated across studies (Fig. 2C). Specifically, DE analyses were conducted independently in 22 datasets, comprising 11 substudies from the HLCA core datasets and 11 datasets from Wauters *et al.* [5]. For each dataset, DE testing was performed using the MAST framework, followed by stringent quality filters (*p_adj_ <* 0.05, *log*_2_*FC* 1). It leads to consensus marker panels, which comprised 689 genes at Level 1, 753 at Level 2, and 1065 at Level 3. Stringent reproducibility thresholds were applied, markers were required to be detected in 10 out of 11 studies for most cell types, with progressively relaxed criteria (3 out of 11 studies) for rarer populations such as mast cells (18 markers) and pDCs (10 markers) (Fig. 2C; Supplementary Fig. S1 to Fig. S6;). The detailed robust marker list per cell type for Level 1 to Level 3 are listed in the Supplementary Table 1. Cell-type-specific markers were further capped at 200 genes when counts exceeded this threshold (e.g., CD4^+^ T cells: 223→200; epithelial: 308→200; ciliated: 505→200). To ensure comprehensive biological representation, additional curated markers were manually appended for neutrophils and pDCs at Level 2, and monocyte, macrophage, and alveolar subtypes at Level 3. After removing the duplicate genes among different cell types, the final list comprised 664 genes at Level 1, 723 at Level 2, and 851 at Level 3.

The number of marker genes identified varied substantially among cell types (Fig. 2D). Epithelial cells, T cells (both CD4^+^ and CD8^+^), and macrophage populations displayed the largest marker repertoires (*>*200) while others such as plasma cells and neutrophils had relatively few (*<*10), reflecting greater transcriptional diversity or stronger within-type expression signatures in the former. These differences underscore variation in marker gene availability across broad versus more specialized immune lineages. These marker panels provide a standardized foundation for consistent cell-type annotation in future BAL or lung single-cell studies. Despite this heterogeneity, the heatmaps of top marker genes across cell types confirmed strong cell-type-specific expression patterns, supporting the reliability of these markers across independent datasets (Fig. 2E). Distinct clusters with minimal cross-type overlap highlight the specificity of the selected markers and the robustness of our harmonized annotation pipeline, providing a reproducible framework for integrating BAL single-cell data across studies.

Overall, the resulting taxonomy and marker catalogues provide (i) a consistent backbone for cross-study BAL harmonization and (ii) a robust, reusable feature space for automated annotation under dataset shift. We release versioned resources (Level 1/2/3 marker lists) (https://github.com/yushannn/BALannotation/tree/main/result), which were used in our downstream process for ensemble modelling, independent validation, and disease comparisons.

### 2.2 Cell type heterogeneity and functional diversity in BAL macrophages at Level 3

While the HLCA provides a comprehensive reference for lung immune and epithelial populations, it does not include annotations at the fine-grained resolution defined in our Level 3 taxonomy. In particular, macrophages required reclustering to uncover additional transcriptionally distinct subsets that were not previously resolved in existing BAL atlases. To characterize this diversity at the finest resolution (Level 3), we performed detailed subclustering and cross-study marker comparison of BAL macrophages (Fig. 3A-B). The UMAP visualization at Level 2 revealed a broad separation of alveolar and non-alveolar macrophages, indicating transcriptionally distinct macrophage compartments in the lung environment (Fig. 3A). These two major populations were further divided into multiple Level 3 subclusters (Fig. 3B), each characterized by unique gene expression signatures (Fig. 3C). Specifically, we identified several transcriptionally stable subtypes, including CCL2^+^, CCL18^+^, RGS1^+^, MT1G^+^ macrophages, as well as smaller subsets associated with inflammatory or tissue-remodeling functions.

**Fig. 3.**
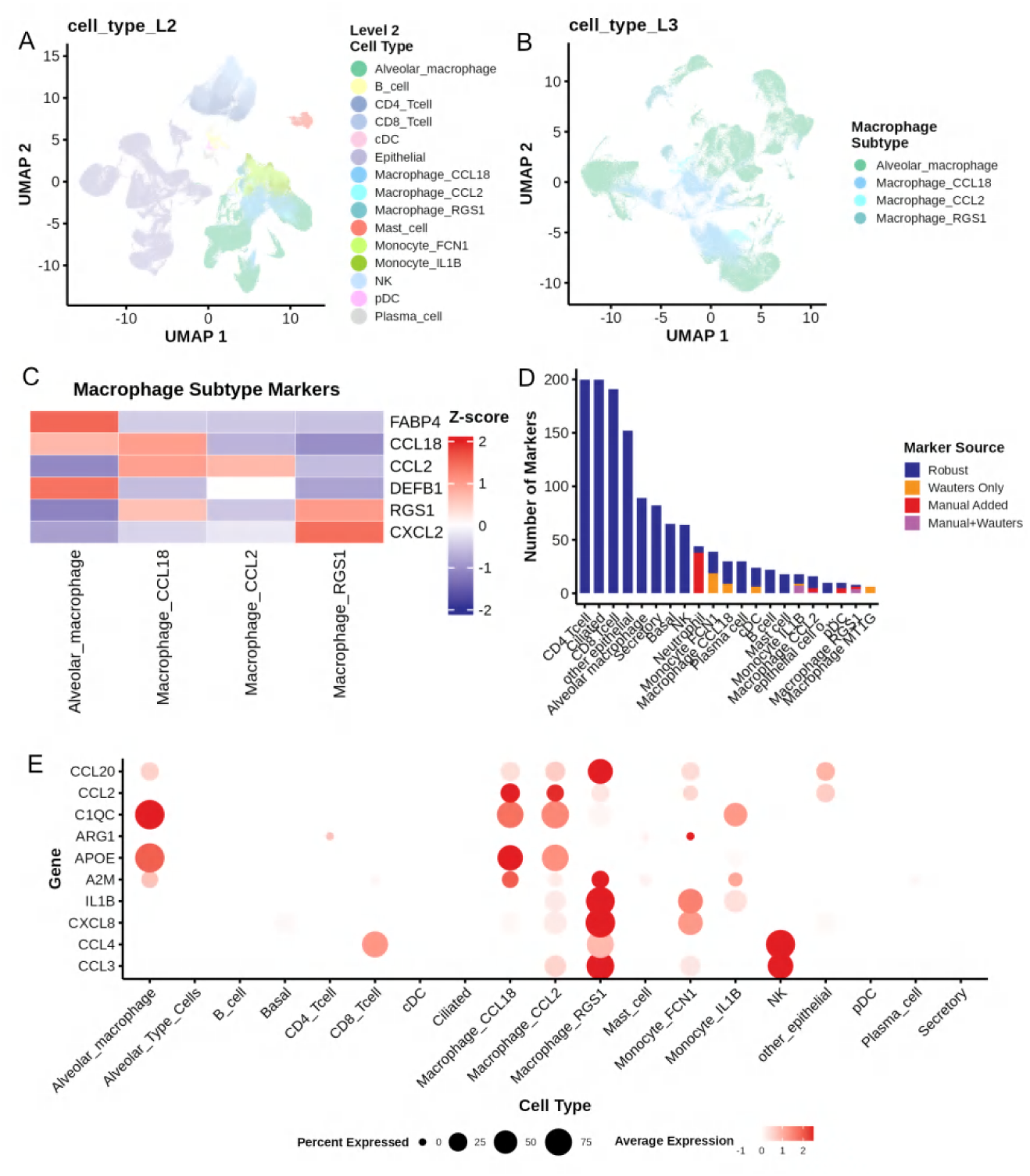
Cell Type Heterogeneity and Functional Diversity in BAL Macrophages at Level 3. (A) UMAP of the HLCA core dataset showing Level 2 cell-type annotations. The projection of 347,333 cells reveals substantial transcriptional heterogeneity within major cell type categories, particularly in macrophage and epithelial populations. (B) Macrophage subtypes: Functional diversity. UMAP of 111,923 macrophage cells colored by Level 3 subtype (alveolar macrophage, macrophage CCL18, macrophage CCL2, macrophage RGS1). Distinct spatial clustering indicates coordinated transcriptional programs associated with different activation states. (C) Marker gene expression across macrophage subtypes. Heatmap showing scaled expression of representative marker genes (FABP4, CCL18, CCL2, DEFB1, RGS1 and CXCL2) across four macrophage subtypes, supporting the biological distinctiveness of Level 3 annotations. (D) Diversity and contribution of marker gene sources for Level 3 cell-type annotation. Stacked bar plot summarizing the origin of marker genes across Level 3 cell types, including Robust, Wauters *et al.* Only, Manual Added, Manual+Wauters *et al.*, illustrating the complementary contribution of manual curation. (E) Specificity of manual marker genes across fine-grained cell types. Dot plot showing expression specificity of the top 10 manually curated marker genes across Level 3 cell types. Dot size represents the fraction of expressing cells and color indicates average expression level, demonstrating the added resolution enabled by manual annotation.

The diversity of marker gene evidence (Fig. 3D) further underscored this robustness: many macrophage markers were supported by both the HLCA and Wauters *et al.* datasets, whereas a smaller fraction was study-specific, reflecting biological or technical variability. Furthermore, dot plots of the top 10 marker genes from Level subtypes (Fig. 3E) illustrated distinct transcriptional programs, with CCL18^+^ and MT1G^+^ macrophages showing features of tissue residency, and CCL2^+^ and RGS1^+^ subsets displaying inflammatory activation. These analyses collectively reveal that BAL macrophages encompass diverse, reproducible transcriptional states not captured in prior reference annotations. In Fig. 3C, no HLCA cluster showed dominant overlap with the MT1G^+^ subtype, suggesting that this stress-responsive state, while evident in BAL, is underrepresented in tissue-derived HLCA macrophages. This is biologically plausible: MT1G^+^ stress-response macrophages are often enriched in lavage but underrepresented in tissue-derived references. These distinct gene expression programs matched the signatures defined in the Wauters’ BAL dataset, providing strong evidence that the reassignment is biologically consistent.

### 2.3 Internal validation performance (LOSO cross-validation)

We assessed model performance using a leave-one-study-out (LOSO) cross-validation scheme on the internal training cohort (HLCA + Wauters; 11 studies, 387,545 cells). Fig. 4A shows the per-fold validation accuracy for each left-out study using linear discriminant analysis (LDA) and random forest (RF) as base learners. Both methods achieved consistently high accuracy across most studies, typically exceeding 75%. For several datasets, including Leroy *et al.*, Budinger *et al.*, Misharin *et al.*, and Meyer *et al.*, performance reached or exceeded 90% for both algorithms, whereas others (Meyer *et al.*, Rojas *et al.*, Lafyatis *et al.*, Kropski *et al.*, and Nawijn*et al.*) showed reduced performance, reflecting reduced performance/intrinsic study heterogeneity, particularly with LDA. While RF displayed slightly greater variability across studies, it generally outperformed or matched LDA. These results suggest that predictive performance is broadly robust across cohorts, though study-specific heterogeneity remains. These internal validation outcomes established the foundation for ensemble integration and external testing.

**Fig. 4.**
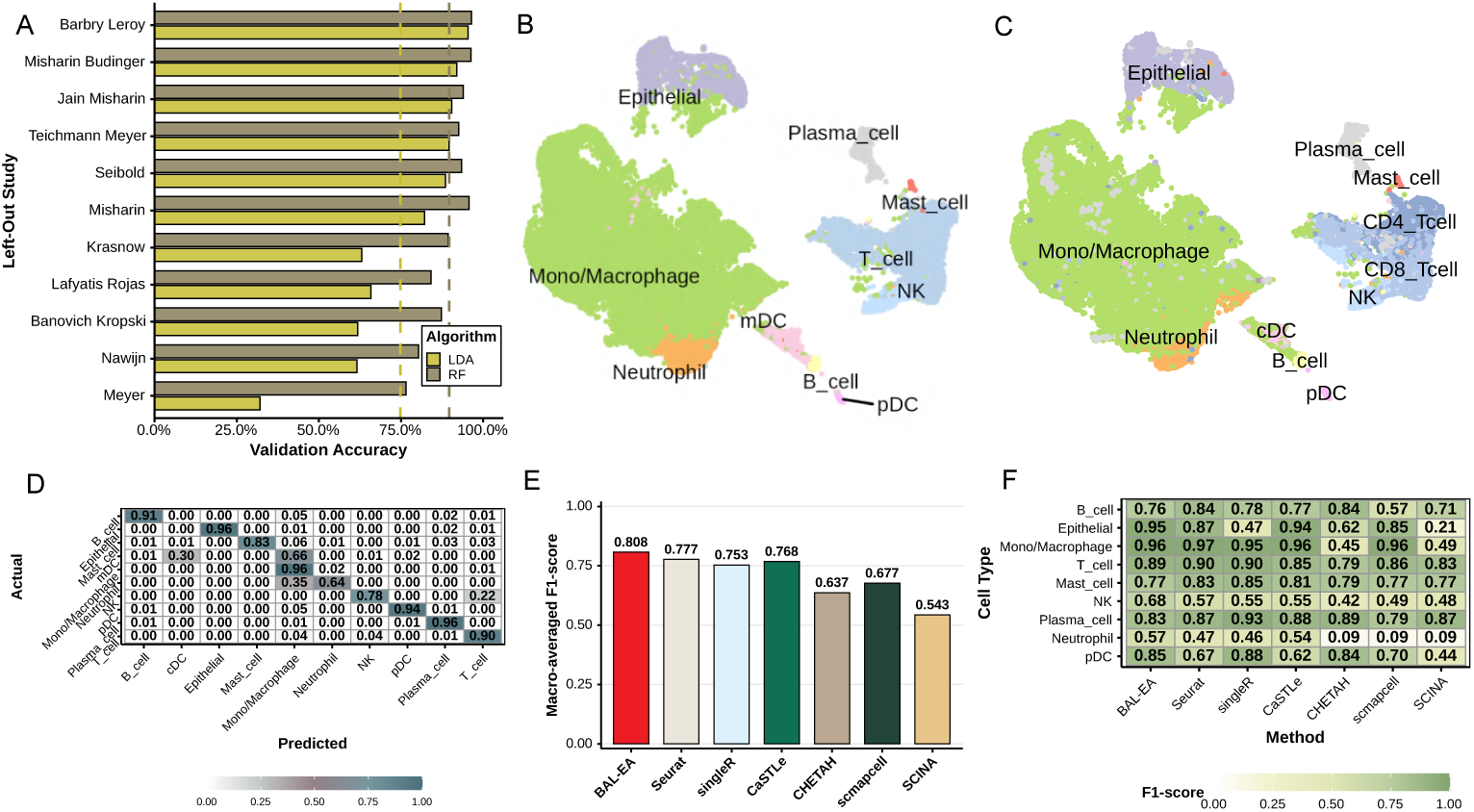
BAL-EA Performance Validation and Benchmarking Against State-of-the-Art Methods. (A) LOSO internal validation accuracy for LDA and RF across 11 studies; each bar represents one study held out for testing. (B) The UMAP plot for the Liao *et al.* dataset. (C) The same UMAP as in (B) but with cell types annotated using the BAL-EA model. (D) Confusion matrix for BAL-EA predictions. Rows represent true cell types, and columns represent predicted types; darker blue indicates higher recall, and off-diagonal entries show misclassifications. (E) Macro-averaged F1 score comparison of BAL-EA (red) with six state-of-the-art methods (Seurat, SingleR, CaSTLe, CHETAH, scmapcell, SCINA) on the external validation dataset (Liao *et al.* dataset). Numbers above bars indicate exact F1 scores. (F) Per-cell-type F1 score heatmap for nine cell types (columns) across seven methods. Darker green indicates higher performance. No clustering applied.

### 2.4 External validation on an independent BAL cohort

To assess generalization under realistic domain shift, we evaluated BAL-EA on an external BAL cohort not used for training: the COVID-19/healthy BAL dataset from Liao *et al.* [4]. Fig. 4B and C show the UMAP plots for cell type annotations at level 1 from the conventional clustering approach reported in Liao *et al.*[4] and from our BALEA predictions, respectively. For UMAP plots with additional resolutions (level 2 and level 3) are shown in Fig. S7. Despite minor differences in naming conventions, BAL-EA predictions closely recapitulated the published clustering structure, as evident from the great alignments of major cell types in both figures, including epithelial cells, mono/macrophages, and T cells. Within mono/macrophage compartments, alveolar macrophages, non-alveolar macrophages, and their subtypes formed well-defined clusters corresponding to the original annotation. Similarly, B cells, plasma cells, T cells, and epithelial cells exhibited consistent localization, underscoring the cross-dataset robustness of BAL-EA label transfer.

Discrepancies were primarily confined to granulocytic and cytotoxic populations. A fraction of neutrophils were predicted as macrophages, and some NK cells were assigned to T cells. The confusion-matrix heatmap (Fig. 4D) quantifies these outcomes. Correctly classified proportions (diagonal entries) reached accuracy 0.96 for mono/macrophages, epithelial cells, and plasma cells, respectively; 0.94 for pDCs; 0.91 for B cells; and 0.90 for T cells. Lower accuracies were observed for NK (0.78) and neutrophils (0.64), the two most error-prone lineages. Most neutrophil errors reflected transcriptional overlap with the monocyte/macrophage axis (35% misassigned), whereas NK-T confusion (~22%) arose from shared cytotoxic markers (PRF1, GZMB). The pDCs remained largely well separated, with only ~5% misclassified as macrophages.

We next benchmarked BAL-EA against six state-of-the-art auto-annotation tools [20] using macro-averaged F1 scores across major cell types. BAL-EA achieved the highest overall macro-F1 (0.808), surpassing Seurat (0.777) [21], SingleR (0.753) [13], CaSTLe (0.768) [15], CHETAH (0.637) [16], scmapcell (0.677) [14], and SCINA (0.543) [17] (Fig. 4E). In particular, BAL-EA demonstrated balanced and consistent performance across cell types. By cell type, BAL-EA led for epithelial cells (0.95), NK cells (0.68), and neutrophil cells (0.57), and was highly competitive for mono/macrophages (0.96) and T cells (0.89). Although Seurat and SingleR achieved comparable results in specific classes (e.g., SingleR for pDCs 0.882 and plasma cells 0.935, Seurat for mono/macrophages 0.966), BAL-EA delivered the most balanced performance across lineages overall (Fig. 4F).

### 2.5 Translational analyses in COPD

A key motivation for a BAL-centric annotation framework is to enable reproducible, disease-relevant comparisons across cohorts. Using standardized labels and the study-wise ensemble, we quantified cell-composition shifts and myeloid programs in COPD, and interpreted the results in the context of prior bronchoalveolar single-cell studies [3–6, 8–10]. To this end, we applied the BAL-EA to our newly generated large in-house BAL scRNA-seq data comprising 241,924 cells from 30 individuals. The UMAP of the dataset with cell types annotated by the BAL-EA model (Fig. 5A) revealed high concordance with the expected cell type distributions, particularly for the major cell types, including macrophages, epithelial cells, lymphocytes (T cells and NK cells), and others. For comparison, the UMAP showing the Seurat [21] clustering annotations of the same dataset was provided in Fig. S8, illustrating consistent overall cell type structure.

**Fig. 5.**
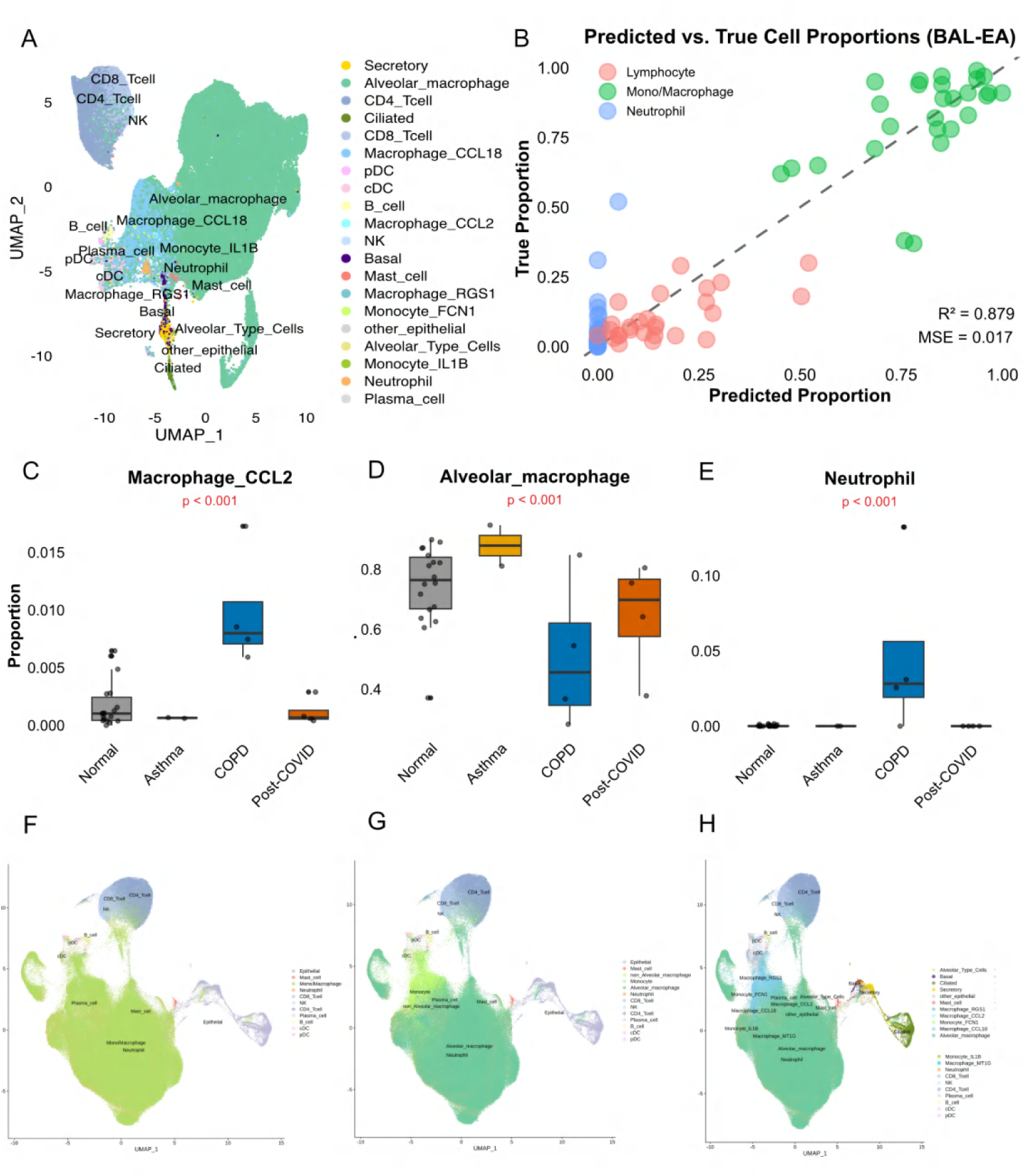
Application on in-house BAL dataset and publicly available BAL datasets. (A) UMAP of the in-house BAL data annotated with the BAL-EA model. (B) Correlation between predicted and measured cell type proportion. The estimated cell type proportion was compared with the true cell type proportion for macrophages, lymphocytes, and neutrophils. (C-E) Differential analysis of cell type proportions between normal and disease samples (COPD, Asthma and Post-COVID). (F-H) UMAPs of integrated BAL single-cell transcriptome atlas at different resolutions.

The proportions of cell types based on the predicted classes per sample were then calculated and compared with the true cell type proportions across macrophages, lymphocytes, and neutrophils. This comparison was performed on 27 of the 30 samples for which the ground-truth cell type proportion measurements were available. High correlations between the estimated proportions and measured proportions were observed (*R*^2^= 0.88) with low mean squared error (MSE = 0.01) (Fig. 5B). When focusing on 15 healthy individuals, the model performance improved substantially, achieving higher correlation (*R*^2^= 0.92) and lower error (MSE = 0.011) refer to Fig. S9A. Compared with other auto-annotation methods, including CHETAH, SCINA, Seurat, CaSTLe, and scmapcell, BAL-EA demonstrated the lowest error for macrophage quantification (MSE=0.047) in Fig. S9B. This is particularly important given that macrophages represent the predominant cell population in BAL samples and exhibit extensive heterogeneity with diverse functional subtypes that serve as key indicators of pulmonary immune status and disease progression. It validated the model’s accuracy in estimating cell type proportions, underscoring the reliability of BAL-EA for quantitative analyses.

Furthermore, cell type proportions of major cells between normal and COPD BAL samples were compared (Fig. 5C-E). For this purpose, previously identified subtypes of mono/macrophage along with other major cell types were used. The analysis revealed significant increases in macrophages CCL2, and neutrophils in COPD patients compared to healthy controls. These findings suggested enhanced inflammation in COPD. In contrast, a notable decrease in AMs was observed in COPD samples (binomial GLM with age and sex adjustment, p-values *<* 0.001 for all three cell types) compared to the normal group, reflecting a potential disruption of homeostatic macrophage populations in the disease state. Additionally, supplementary analysis showed significant increases in pro-inflammatory subsets, including macrophages CCL18, macrophages RGS1, and monocytes FCN1, in COPD patients (p-values *<* 0.001, in Fig. S10), further supporting the inflammatory phenotype observed in this disease.

### 2.6 The BAL single-cell atlas and public resource

We release a versioned resource that unifies the BAL-centric taxonomy and cross-study markers, the study-wise ensemble classifiers and baselines, external validation artifacts, and disease-focused summaries. The atlas includes annotated objects at three resolutions (Levels 1 to 3), consensus marker catalogues with reproducibility tags, pre-trained per-study LDA models and the stacked ensemble, as well as evaluation outputs for the external cohort of Liao *et al.* [4]. Rather than performing clustering on the integrative dataset from scratch, which needs intensive batch effect removal and manual adjustments of resolutions, BAL-EA automatic cell type annotation facilitates quick and robust integration. To this end, seven publicly available BAL scRNA-seq datasets, together with the in-house BAL scRNA-seq dataset were collected [4–10]. Each BAL dataset was processed and re-annotated using the BAL-EA with three cell-type resolutions (Fig. 5F-H).

## 3 Discussion

In this study, we present a comprehensive framework for automated cell type annotation for human BAL scRNA-seq data. Unlike existing general-purpose annotation tools, our BAL-EA model leverages BAL-specific marker genes and integrates prior knowledge from the HLCA, resulting in improved annotation accuracy. Notably, this framework is flexible on training models with different cell type resolutions. Moreover, the models can be easily updated as new scRNA-seq datasets become available, ensuring their continued relevance as the field progresses. In addition, by leveraging the BAL-EA model and integrating the largest inhouse lung BAL scRNA-seq data, we constructed the most comprehensive BAL single-cell transcriptomic atlas to date, encompassing both healthy and diseased states. In summary, this study not only provided significant advancements in the study of BAL samples in both health and respiratory diseases but also offers an effective framework for creating automatic cell type annotation for other tissues.

Cell type annotation is a critical prerequisite but also a key challenge in scRNA-seq analysis. Traditional clustering-based strategies involve extensive manual interventions, introducing bias and inconsistency in the resolution, specificity, and even naming conventions of identified cell populations [22, 23]. It further impacts reproducibility and scalability across larger, diverse datasets. The creation of a cell atlas offers a coherent and unbiased reference for all cell types, significantly facilitating the annotation of new samples. However, integrating diverse datasets and performing clustering can be both time-consuming and computationally demanding. For instance, clustering millions of cells often requires high-performance computing resources with extremely large memory capacity, which may not be readily available in research labs. Despite these challenges, such rich resources provide an invaluable foundation for building automated annotation models. We hypothesize that an automatic annotation strategy will streamline this process, or at least to a certain extent alleviate the computational burden, with the advent of a robust and reliable reference dataset. While no specific BAL cell atlas currently exists, the HLCA [12] is available, though its application to human BAL cells has not been fully validated. In this study, we explored the potential of developing a BAL-specific cell atlas using a machine-learning-based autoannotation strategy. This was implemented by integrating BAL scRNA-seq datasets with the HLCA dataset and by utilizing an ensemble-learning approach. Through ensemble learning, the large-scale HLCA dataset was harmonized with the limited but high-quality BAL scRNA-seq datasets.

Another major achievement of this study is the identification of cell-type-specific robust marker genes at various resolutions, facilitating accurate and nuanced annotation of BAL cell populations. These marker genes provide a foundational resource for distinguishing between major cell types (e.g., macrophages, T cells, and epithelial cells) as well as their subtypes (e.g., AMs, monocyte-derived macrophages, CD4 and CD8 T cells). This stratification enables precise exploration of cellular heterogeneity and offers insights into functional diversity within BAL samples, even under pathological conditions. Furthermore, the robust marker genes might provide a powerful basis for accurate bulk RNA-seq data cell type deconvolution.

Peer comparisons with other annotation tools and clinical applications of our model further validated the effectiveness of the BAL-EA framework. In benchmarking analyses, the BAL-EA model consistently outperformed existing tools, particularly for challenging cell populations such as neutrophils and NK cells. Additionally, its application to clinical datasets demonstrated its ability to uncover biologically meaningful differences in cell type proportions, such as increased neutrophils and macrophage CCL2 subtypes and decreased AMs in COPD patients.

Despite several advances, there are limitations to consider. Our analysis strategy aims to implement an automatic cell type annotation. The resolution or the number of cell types as well as good-quality reference datasets should be provided before building the automatic prediction models, which limits the efficiency of automatic annotation for clinical applications. However, similar to the clustering-based annotation where specific resolution needs to be defined before determining the detailed cell types, it would be straightforward to train automatic annotation models at different resolutions. It is possible that certain cell types in new samples, particularly those from diseased states or representing rare populations, may not be included in the existing BAL atlas. In the scenario of a new cell type emerging, we are committed to updating the models promptly to ensure our model or tool remains accurate and up-to-date.

## 4 Methods

### 4.1 Cohorts and datasets

#### 4.1.1 Training datasets

**HLCA 2023** The HLCA core was constructed from 584,444 high-quality scRNA-seq profiles derived from 166 lung tissue samples of 107 healthy individuals across 11 studies[12]. Datasets were integrated using scANVI [24], which corrects batch effects while preserving biological variation. The HLCA core features a consensus annotation of 61 cell types, established through re-annotation combining harmonized original labels and expert consensus. This HLCA core atlas captures substantial cellular and biological diversity across demographics, experimental protocols, and anatomical locations along the proximal-to-distal axis of the respiratory system.

The HLCA has gone through complete preprocessing and curation by their consortia, which included rigorous quality control and removal of low-quality cells. Ambient RNA contamination and doublet artifacts were either explicitly addressed in the pre-processing pipelines or indirectly minimized during quality filtering and integration across studies. Because of this, the released objects are considered “analysis-ready” and suitable for use as reference atlases for label transfer or comparative analyses. Re-running ambient correction or doublet detection on these processed objects would risk over-filtering and discarding biologically relevant cells and could introduce inconsistencies with the published consensus annotations.

**Wauters *et al.* 2021** BALF scRNA-seq reference data were obtained from Wauters *et al.* [5], who performed comprehensive immune profiling of BALF samples from 24 patients with mild or critical COVID-19 and 13 healthy controls. Using the 10x Genomics platform, they generated 65,166 high-quality single-cell transcriptomes with detailed annotation and pseudotime analysis of major immune and epithelial compartments. The dataset includes diverse lymphoid (CD4+ and CD8+ T cells, NK cells, B cells, and plasma cells), myeloid (monocytes, alveolar and monocyte-derived macrophages, neutrophils, and dendritic cells), and epithelial populations, capturing the phenotypic and transcriptional diversity of the lung immune microenvironment across disease conditions. The raw unique molecular identifier (UMI) read count data can be downloaded from https://lambrechtslab.sites.vib.be/en/immune-atlas.

Following the procedure proposed in Wauters *et al.* [5], UMI read counts were normalized using CPM-style scaling and subjected to stringent quality control filtering (e.g., 200–5000 detected genes per cell and *<*20% mitochondrial reads), while genes expressed in fewer than 3 cells were excluded. UMI counts were normalized using counts per million (CPM) transformation followed by log transformation: log(CPM + 1). To minimize disease-related confounding effects, analyses were restricted to healthy control samples only. The processed dataset was then partitioned by patient identifier, with samples randomly assigned across 11 cross-validation folds (10 samples per fold) to facilitate subsequent model training and validation.

#### 4.1.2 Validation dataset

**Liao *et al.* 2020** BALF scRNA-seq dataset (GSE145926), published by Liao *et al.* [4], provides a comprehensive transcriptomic reference of bronchoalveolar immune cells from COVID-19 patients and healthy controls, including three patients with moderate COVID-19, six patients with severe or critical COVID-19, and three healthy controls, with an additional healthy BALF sample incorporated from a public source. scRNA-seq libraries were prepared using the 10x Genomics platform, and the libraries were sequenced on the BGI MGISEQ-500. Data processing included stringent quality control, UMI filtering, cell clustering, and batch integration using Cell Ranger and Seurat. In total, 66,452 cells were analyzed, yielding a landscape of pulmonary immune cell heterogeneity defined by major lineages such as macrophages, neutrophils, dendritic cells, NK cells, T cells, B cells, plasma cells, and epithelial cells. Cell type annotation was supported by canonical marker gene expression and supported by clustering and downstream pathway analysis. The processed data used in this study were directly downloaded http://cells.ucsc.edu/covid19-balf/nCoV.rds as the Seurat object.

#### 4.1.3 Application datasets

**Morse *et al.* 2019** The GSE128033 dataset includes two healthy human BALF scRNA-seq samples profiled by Morse *et al.* [19] using 10x Genomics and sequenced on an Illumina NextSeq 500 with the 3’ gene expression (V2) chemistry. After standard quality control and filtering, UMI or expression matrices were generated, resolving multiple pulmonary immune cell types, including distinct FABP4^+^, SPP1^+^, and FCN1^+^ macrophages, as well as lymphocyte subpopulations. In total, 5,126 BAL cells and 23 cell types were obtained. However, no publicly available cell-type annotation file was found for this dataset.

**Chua *et al.* 2020** The Chua *et al.* [8] dataset contains scRNA-seq profiles from nasopharyngeal and BALF of patients with moderate or critical COVID-19 as well as from healthy controls (eight moderate, eleven critical, and five healthy controls). Among these samples, only two BAL samples were derived from COVID-19 patients. The two BAL samples were processed using the 10x Genomics Single Cell 3’ Library platform and sequenced on an Illumina NovaSeq 6000. Data were analyzed with CellRanger and Seurat following standard quality control and normalization. Cell type annotations were guided by canonical marker genes and clustering results. Raw reads data are controlled accessible via the European Genome-phenome Archive (EGA, accession id EGAS00001004481, https://ega-archive.org), and processed count matrices and cell metadata (including cell type, sample, case severity, and quality metrics) can be directly downloaded from FigShare (https://doi.org/10.6084/m9.figshare.12436517). After QC, 38,658 BAL cells were retained for downstream analysis.

**Mould *et al.* 2021** The BALF scRNA-seq dataset GSE151928, published by Mould *et al.* [10], was generated from BALF samples of 10 healthy adults collected via fiberoptic bronchoscopy. Single-cell suspensions were captured using the 10x Genomics, and the libraries were sequenced on the Illumina NovaSeq 6000. The raw UMI count data are available in GEO under the accession number GSE151928.

Following the Seurat pipeline for scRNA-seq data processing and quality control, the final dataset consisted of 16,251 cells and 3,816 genes. Analyses identified a diverse landscape of immune cell types, including resident macrophages, monocyte-like cells, dendritic cells, and lymphocytes, with distinct alveolar macrophage (AM) subsets showing pro-inflammatory or metallothionein-related signatures and rare matrix-associated monocytes.

**Grant *et al.* 2021** The dataset GSE155249, published by Grant *et al.* [6], comprises BALF scRNA-seq profiles of 10 patients with severe COVID-19 within 48 hours of intubation. Following flow sorting of macrophages and T cells, single-cell libraries were prepared using the 10x Genomics 5’ platform and sequenced on an Illumina NovaSeq 6000. Processed gene expression matrices were generated with Cell Ranger and integrated using Scanpy [25]. Doublets were detected and removed with scrublet [26], ambient RNA was corrected with FastCAR, and multisample integration was performed with BBKNN. The final quality-controlled dataset consisted of 105,715 cells.

**Li *et al.* 2022** The GSE193782 dataset, published by Li *et al.* [9], provides BALF scRNA-seq profiles of four healthy control subjects and three individuals with mild, uninflamed cystic fibrosis (CF). The study profiled 113,213 cells using the 10X Genomics platform, followed by rigorous data processing and integration in Seurat. It investigated the cellular and transcriptional profiles between CF patients with normal lung function and minimal lung inflammation compared with healthy controls. Unbiased clustering and thorough marker gene analysis revealed twelve major cell types, including AM, monocyte subtypes, dendritic cells, and epithelial cells, with special emphasis on AM heterogeneity.

**He *et al.* 2023** The scRNA-seq dataset GSE147143 [7] contains 4,264 high-quality transcriptomic profiles from BALF of three patients with severe COVID-19. Libraries were prepared using the 10x Genomics platform and sequenced on an Illumina HiSeq X Ten. It profiled leukocytes and epithelial cells to investigate the cellular and molecular mechanisms underlying acute respiratory distress syndrome (ARDS). The dataset includes annotations for major pulmonary cell populations, such as epithelial cells (club, ciliated, AT1, and AT2 cells), endothelial cells, and immune cells, including macrophages, monocytes, neutrophils, B cells, and NK & T cells, although a publicly available annotation metadata file was not found.

**The inhouse dataset** comprises BAL scRNA-seq data from 30 samples, of which 27 have clinical metadata including 15 healthy controls and patients with COPD (n=4), post-COVID-19 (n=5), asthma (n=2), and bronchiectasis (n=1). The single-cell capture was completed using the Chromium Next GEM Single-Cell 3’ Kit v3.1 and Chip G (10x Genomics, CA, USA). Libraries were prepared according to the 10X Genomics Chromium Single-Cell 3’ Reagent Kits User Guide. The final libraries sequencing was performed on the NextSeq 2000 (Illumina, CA, USA), targeting the sequencing depth of 40,000 reads per cell. Raw reads (fastq files from each sequencing run) were processed via Cell Ranger (version 8.0.0) using the human (Homo sapiens) reference genome (GRCh38.p14, GENCODE release 46) for alignment and UMI quantification. The R package Seurat was used to preprocess the scRNA-seq data. Cells with *<*200 or *>*10,000 detected genes (nFeature RNA), mitochondrial gene content *>*10%, or total molecule counts (nCount RNA) *>*1,000 were excluded from downstream analyses. After preprocessing and quality control, a total of 241,924 high-quality cells were collected for downstream analysis.

**Table 1.** Overview of the datasets used during this study

| Dataset | Cells Num | Genes Num | Year | CT | Description | Platform | Data source |
| --- | --- | --- | --- | --- | --- | --- | --- |
| In-house | 241,924 | 33,465 | – | – | Human lung BAL | 10X | – |
| HLCACore<sup>a</sup> | 584,944 | 28,024 | 2023 | 13 | Human healthy lung tissue | – | Human Cell Atlas Data Portal [12] |
| Li<sup>a,b</sup> | 113,213 | 27,805 | 2022 | 12 | Human healthy and CF BAL | 10X | GSE193782[9] |
| Grant | 105,715 | 21,819 | 2021 | 28 | Human lung ICU | 10X | GSE155249[6] |
| Wauters<sup>b</sup> | 65,166 | 33,538 | 2021 | 18 | Human healthy and COVID-19 BAL | 10X | Lambrechts lab site[5] |
| He<sup>b</sup> | 4,264 | 21,701 | 2020 | 10 | Human COVID-19 BAL | 10X | GSE147143[7] |
| Mould<sup>a,b</sup> | 16,251 | 3,816 | 2020 | 6 | Human healthy lung BAL | 10X | GSE151928[10] |
| Liao<sup>a,b</sup> | 51,631 | 23,916 | 2020 | 10 | Human healthy and COVID-19 lung BAL | 10X | GSE145926[4] |
| Chua<sup>a</sup> | 38,658 | 25,000 | 2020 | 24 | Human lung disease BAL | 10X | figshare[8] |
| Morse | 1,200 | 11,852 | 2019 | 16 | Human healthy lung BAL | 10X | GSE128033[19] |

<sup>a</sup>Raw data download available; <sup>b</sup>Only include BAL samples; CT: the number of cell types.

### 4.2 In-house data collection and processing

A full description of the bronchoscopy and sample procurement has been described elsewhere [2]. In brief, to obtain BAL, a bronchoscope was wedged in the lingular segment of the left lung or the right middle lobe. Next, 200 ml of saline was instilled through the suction channel aiming to retrieve at least 30 ml of BAL [2]. Post collection, the BAL processing involved filtration using a 70 micron strainers (VWR, Radnor, Pennsylvania), followed by multiple centrifugations at 300 g, 4°C for 10 minutes and resuspension with 10 ml Dulbecco’s phosphate-buffered saline (DPBS) (Gibco, Waltham, Massachusetts, USA) with 0.4% fetal bovine serum (FBS) (Invitrogen, Waltham, MA, USA) [2]. The resultant single cell suspension was then transferred to the UBC Sequencing Facility at the Biomedical Research Centre (BRC-seq, University of British Columbia) for capture and sequencing.

### 4.3 Label harmonization and BAL-centric taxonomy

To standardize cell-type labels across BAL scRNA-seq datasets, we implemented a hierarchical mapping strategy. Original cluster identities from each study were manually mapped to three consensus levels using curated marker genes and previously published annotations [4, 5, 12].

Level 1 consolidated broad immune and epithelial compartments (n=11 classes), including B cells, CD4^+^ T cells, CD8^+^ T cells, cDCs, epithelial cells, mono/-macrophage, mast cells, natural killer (NK) cells, neutrophils, plasma cells, and pDCs. Clusters annotated as “Mono/Macrophage” are defined to include the mononuclear phagocyte lineage, comprising alveolar macrophages, non-alveolar macrophages, and monocytes. The term was chosen because these populations constitute a transcriptional continuum in BAL, especially in inflammatory or remodeling pulmonary environments where monocyte-to-macrophage differentiation is occurring. The collective designation at Level 1 reflects their common biological function as innate immune phagocytes and is suitable for general categorization in cross-dataset evaluations. Level 2 refined myeloid heterogeneity by distinguishing alveolar macrophages, non-alveolar macrophages, and monocytes (n=13 classes). Other lineages remained grouped as in Level 1. Level 3 provided fine-grained resolution (n = 21 classes), incorporating transcriptionally distinct monocyte subsets (FCN1^+^, IL1B^+^, HSPA6^+^) and non-alveolar macrophage programs (CCL2^+^, CCL18^+^, RGS1^+^, MT1G^+^). Epithelial cells were partitioned into basal, ciliated, secretory, alveolar epithelial, and other epithelial cells. These mappings were formalized through R functions to ensure reproducibility and consistent application across datasets.

Although the HLCA is a standardized lung-wide ontology, its tissue-centric framework inadequately represents the composition of BALF. Directly embedding BAL cells into the HLCA space may misclassify lavage-derived populations into parenchymal compartments (e.g., stromal or endothelial cells) that are absent from BAL, whereas HLCA subtypes often span a broader range than those present in lavage samples. In contrast, crucial BAL-enriched lineages, such as neutrophils, are often underrepresented or absent. To preserve biological fidelity while enabling cross-study comparability, we implemented a BAL-centric taxonomy. At coarser resolutions (level 1, n = 11 classes; level 2, n = 13 classes), manual curation was applied to reconcile naming inconsistencies and to unify related subsets across the HLCA and BAL datasets, without performing additional re-clustering. At the finer resolution (level 3), reproducibility-guided marker identification was applied to resolve subtypes. Since HLCA level 3 annotations did not sufficiently capture non-alveolar macrophage heterogeneity, we refined these clusters by integrating subtype-specific transcriptional programs derived from Wauters *et al.* Overlap analyses between Wauters’ signatures and HLCA macrophage clusters enabled one-to-one mapping, which was used to update macrophage subtype annotations. This framework aligns with HLCA at general levels while offering finer resolution tailored to BAL biology. This hierarchical system yields interpretable and reproducible annotations optimized for BAL-specific analyses.

### 4.4 Cross-study marker discovery

As an initial step, common cell types between the HLCA core and the Wauters BAL dataset [5] are processed and manually mapped to establish cross-dataset comparability. For each resolution level (Levels 1–3), cluster labels were harmonized using the R mapping functions. DE analyses were performed independently within each HLCA v1.0 core sub-study (n = 11) and within resampled control subsets of the Wauters BAL dataset to avoid batch-related artifacts. For each study, cell-type-specific differentially expressed genes (DEGs) were identified using Seurat’s FindAllMarkers function (test.use = “MAST”), retaining genes with an adjusted p-value *<* 0.05 and log_2_ *FC* 1. For the Wauters’ dataset, a bootstrap-style resampling was performed by randomly holding out three of the 13 samples as a test set and using the remaining 10 for training. This procedure was repeated 11 times, corresponding to the number of HLCA sub-studies, and the resulting DEG lists from each iteration were integrated with those from the respective HLCA analyses. Marker genes were compared across studies to identify reproducible, cross-study signatures that generalize across independent datasets. Specifically, DE analysis results from the 11 HLCA sub-studies and the 11 Wauters resampling rounds were then aggregated to derive a unified set of robust cell-type-specific markers. For each cell type, DEGs were ranked by reproducibility across datasets, and a threshold was applied to retain genes detected in multiple independent analyses. Thresholds were determined hierarchically, starting from genes recurrent in ≥10 datasets and relaxed stepwise (9, 8, . . ., 3) until each cell type yielded a usable marker panel (*>*5 genes). This adaptive filtering ensured that abundant lineages met stringent consensus criteria, while rarer populations were still represented by reproducible markers. The final consensus marker panels were compiled into the supplementary files for each resolution level. The curated HLCA core served as the reference framework for consistent annotation and robust marker validation.

### 4.5 Mapping macrophage programs from Wauters to HLCA

To harmonize macrophage annotations between BAL and lung tissue, the fine-grained Wauters’ macrophage taxonomy was transferred onto HLCA macrophage populations. HLCA cells labelled as non-classical or classical monocytes, alveolar, elicited, or lung macrophages were subsetted and reclustered. Prior to subtype assignment, UMAP visualization of unsupervised clusters was used to assess intrinsic heterogeneity independent of Wauters-derived labels. Subtype-specific signatures were then derived from the Wauters’ dataset by selecting the top 500 DEGs per macrophage cluster, representing four BAL-enriched states: Macrophage CCL18, Macrophage CCL2, Macrophage RGS1, and Macrophage MT1G.

HLCA cells annotated as elicited or lung macrophages were reclustered across multiple parameter settings. For each HLCA cluster, the top 500 DEGs were identified and compared with Wauters’ subtype signatures to quantify gene-set overlap, generating a signature-overlap matrix. HLCA clusters were assigned to Wauters subtypes based on maximum overlap, with a minimum of 30 shared genes; clusters not meeting this threshold were flagged for manual review. Assignments were validated by canonical marker expression (CCL18, SPP1/APOE, RGS1/CXCL2, MT1G/HMOX1) and visualization in UMAP and t-SNE space. Final subtype labels were recorded in the level 3 annotation of the HLCA object for downstream analyses.

### 4.6 Validation of HLCA macrophage subtypes against Wauters’ dataset

#### 4.6.1 Clustering and subtype assignment

HLCA macrophage cells were subjected to unsupervised clustering using Seurat’s FindClusters function with resolution 0.2, based on the top 2,000 variable features and 20 principal components. This resolution was selected to identify distinct functional subtypes while avoiding over-fragmentation. To align HLCA macrophage populations with a BAL-centric taxonomy, we leveraged lineage-specific gene signatures defined by Wauters *et al.* from healthy BAL controls, which capture distinct macrophage activation programs under multiple conditions. HLCA clusters were compared to these Wauters-derived macrophage subtypes (CCL18^+^, CCL2^+^, RGS1^+^) to assign each cluster to the most similar transcriptional program based on gene signature overlap.

#### 4.6.2 DEG analysis and cross-dataset correspondence

For each HLCA cluster, DE analysis was performed using the MAST test (adjusted p-value *<* 0.05, *log*_2_*FC* 1). The resulting DEG lists were compared with marker genes defining the Wauters macrophage subtypes to quantify transcriptional correspondence. Overlaps were visualized as a heatmap, with the number of overlapping genes indicating the degree of correspondence between HLCA clusters and Wauters-defined subtypes. The overlap-driven mapping was used to assess the degree to which HLCA-derived macrophage clusters recapitulate the functional subtypes identified in the independent Wauters’ BAL dataset. This cross-dataset validation supports the robustness and generalizability of the HLCA Level 3 macrophage subtype definitions and enables cross-compartment comparisons between tissue-derived macrophages and those sampled from BALF.

### 4.7 BAL-EA: Ensemble learning via meta-classification

#### 4.7.1 Leave-One-Study-Out base model training

Base model training employed a rigorous LOSO cross-validation strategy to ensure generalizability across heterogeneous single-cell datasets. The training cohort comprised 11 independent studies from the HLCA atlas plus healthy control samples from Wauters *et al.*, totalling 400,894 cells across 18 distinct cell types (see Section 4.3 for annotation harmonization).

For each LOSO fold (*i* = 1 *. . .* 11), the procedure was as follows: (1) Training set — ten HLCA studies (excluding study i) were combined with 10 of the 13 healthy control samples from the Wauters’ dataset to maximize training diversity; (2) Validation set — study i plus the remaining three Wauters’ healthy samples were held out to evaluate model generalization to unseen batches; and (3) Test set — the independent Liao COVID-19 BALF dataset (n = 66,452 cells) was reserved exclusively for external evaluation and not used in any training or validation step.

For each fold, two independent classifiers were trained separately: LDA and RF (500 trees, default mtry). LDA captures linear decision boundaries, while RF models complex non-linear patterns, providing complementary, biologically interpretable predictions. Features were restricted to HLCA-derived DE markers (with optional additional markers for level 3 fine-type annotation) to ensure computational efficiency.

#### 4.7.2 Meta-Learning with XGBoost

Rather than employing fixed weighting schemes or simple majority voting, BAL-EA learns optimal model combination rules through gradient boosting. The core innovation lies in treating base model predictions as meta-features for a secondary classifier that adaptively integrates complementary information across models.

Meta-feature extraction: For each LOSO fold i, we constructed a meta-training dataset where each cell was represented by the soft predictions (probabilities) from the models trained on the remaining 10 folds. This leave-one-study-out strategy at the meta-level prevents information leakage, ensuring the meta-model never observes predictions from models trained on overlapping data.

XGBoost architecture: Multi-class XGBoost (v1.6.0, objective=“multi:softmax”) was used to map meta-features to cell type labels. Critical hyperparameters were nrounds=2 (to prevent overfitting in the low-dimensional meta-feature space), num class=11/13/21, and the default learning rate (*η* = 0.3). The choice of extremely conservative boosting depth was motivated by two considerations: (1) the meta-feature space is substantially lower than typical gene expression space, increasing overfitting risk, and (2) base models already capture complex patterns, requiring only shallow refinement at the meta-level.

Model ensemble: The LOSO procedure yielded 11 independent XGBoost meta-models, each trained on different base model combinations. For the final prediction on the Liao 2020 set, we applied all 11 meta-models and aggregated their predictions via majority voting. This ensemble-of-ensembles strategy provides robustness against model-specific biases and leverages the diversity introduced by LOSO partitioning.

### 4.8 Model evaluation

#### 4.8.1 Evaluation metrics for classification validation

Model performance was quantified using per-class recall and macro-averaged F1 score, widely used metrics for evaluating both cell type-specific and overall annotation accuracy in single-cell classification tasks.

##### Per-class recall (confusion-matrix accuracy)

For each cell type *i*, the proportion of correctly predicted cells was computed as follows:

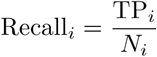

where TP*_i_* denotes the number of correctly classified cells of type *i*, and *N_i_* is the total number of cells belonging to that class. These values correspond to the diagonal entries of the confusion matrix, which summarizes classification performance by comparing predicted versus actual cell type labels. Thus, *Recall_i_* represent cell type–specific accuracies for each population.

##### Precision and F1 score

Precision and recall were combined to calculate the per-class F1 score:

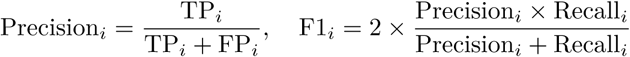

where FP*_i_* is the number of false-positive predictions for cell type *i*. The F1 score reflects the harmonic mean between precision and recall, providing a balanced assessment of classification performance.

##### Macro-averaged F1 score

Overall model performance was summarized by the macro-averaged F1 score:

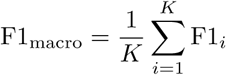

where *K* is the total number of annotated cell types. This metric assigns equal weight to each class, ensuring that both abundant and rare cell type contribute equally to the final score. Macro-F1 values were used to compare BAL-EA with other auto-annotation methods.

#### 4.8.2 Evaluation metrics for clinical validation

For clinical validation analyses, model performance was evaluated by comparing predicted and measured cell type proportions for each cell type across disease and control samples, using regression-based metrics to quantify the agreement, along with statistical tests to assess the significance of differences while adjusting for potential confounders.

##### Coefficient of determination (**R*^2^***)

The goodness of fit between predicted (*y*^*_i_*) and measured or observed (*y_i_*) values was measured using the coefficient of determination:

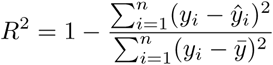

where *n* is the number of samples and *ȳ* is the mean of observed values. *R*^2^ quantifies the proportion of variance in the observed data explained by the model, with higher values indicating better predictive performance.

##### Mean squared error (MSE) and root mean squared error (RMSE)

The overall deviation between predicted and observed values was measured by:

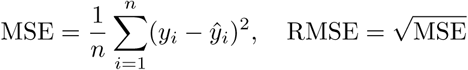

MSE provides an average squared error, while RMSE expresses the same measure in the original units of the target variable, facilitating direct interpretability.

##### Binomial generalized linear model (GLM)

To assess whether cell type proportions differed significantly between disease groups (e.g., COPD vs. control) while adjusting for potential confounders, we applied binomial generalized linear models. For each cell type, cell counts were modeled as

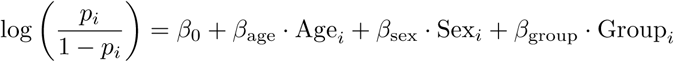

where *p_i_*is the proportion of the cell type in sample *i*, and the model was fitted using the binomial family with the logit link function. The coefficient *β*_group_ represents the log-odds ratio of the cell type proportion between disease and control groups, adjusted for age and sex. Statistical significance was assessed using Wald tests, with a two-sided *p*-value *<* 0.05 considered significant. This approach accounts for the compositional nature of cell type data and controls for demographic covariates that may influence immune cell composition.

### 4.9 Software, versions, and compute environment

All analyses were performed in R (version 4.0 or higher; production runs used R 4.1.2 on high-performance computing servers). Key R packages included Seurat for single-cell data preprocessing and integration; Matrix, MASS, and random Forest for linear discriminant analysis and ensemble classification; xgBoost for meta-ensemble weight learning; dplyr and tidyr for data manipulation; and ggplot2, ComplexHeatmap, cowplot, patchwork, pheatmap, and RColorBrewer for visualization. Complete analysis scripts, intermediate data objects, and figure generation code are available in the BALannotation Git repository at [https://github.com/yushannn/BALannotation. git].

## Supplementary information

Supplementary Table 1. Robust marker gene list per cell type for BAL scRNA-seq at multiple-layers.

Supplementary Materials. Supplementary Figure S1-S10.

## Supporting information

Supplementary Table 1. Robust marker gene list per cell type for BAL scRNA-seq at multiple-layers.

## Acknowledgements.

We acknowledge the Human Lung Cell Atlas consortium for providing harmonized reference annotations and the authors of publicly available BAL datasets that enabled this analysis. This work is supported in part by funds from Genome BC Sector Innovation Program (X.Z., D.D.S), NRC Digital Health and Geospatial Analytics Program (X.Z., X.S., Z.L.), and the Canada Research Chairs (CRC-2021-00232 X.Z.), Michael Smith Foundation for Health Research (SCH-2022-2553 X.Z.), and Mitacs (Y.H., X.Z., D.D.S). Computational resources were provided by the Digital Research Alliance of Canada (the Alliance).

## Data and code availability

- **Data:** The in-house BAL scRNA-seq data generated in this study will be deposited in the Gene Expression Omnibus (GEO). All other single-cell RNA-seq datasets analyzed in this study were obtained from publicly available repositories as indicated in the Method section. Processed data matrices used for benchmarking and model validation are available upon reasonable request to the lead contact.
- **Code:** The BAL-EA framework, along with reproducible analysis scripts and supplementary notebooks, is available at https://github.com/yushannn/BALannotation.git (to be released upon publication).

## Author contributions

Y.H.: Writing – original draft, Writing – Review & editing, Visualization, Software, Methodology, Formal analysis, Investigation, Data curation. Z.L.: Writing – Review & editing, Funding acquisition, Data curation. K.B.: Software, Writing – Review & editing. B.M.: Software, Writing – Review & editing. J.M.L: Sample collection, Writing – Review & editing. F.V.G: Experiment and data acquisition, Writing – Review & editing. X.S.: Study conceptualization, Writing – review & editing, Supervision, Project administration, Methodology, Funding acquisition. X.Z: Study conceptualization, Writing – Review & editing, Supervision, Project administration, Methodology, Funding acquisition. D.D.S: Study conceptualization, Funding acquisition, Writing – Review & editing.

## Appendix A Supplementary Figures

**Fig. S1.**
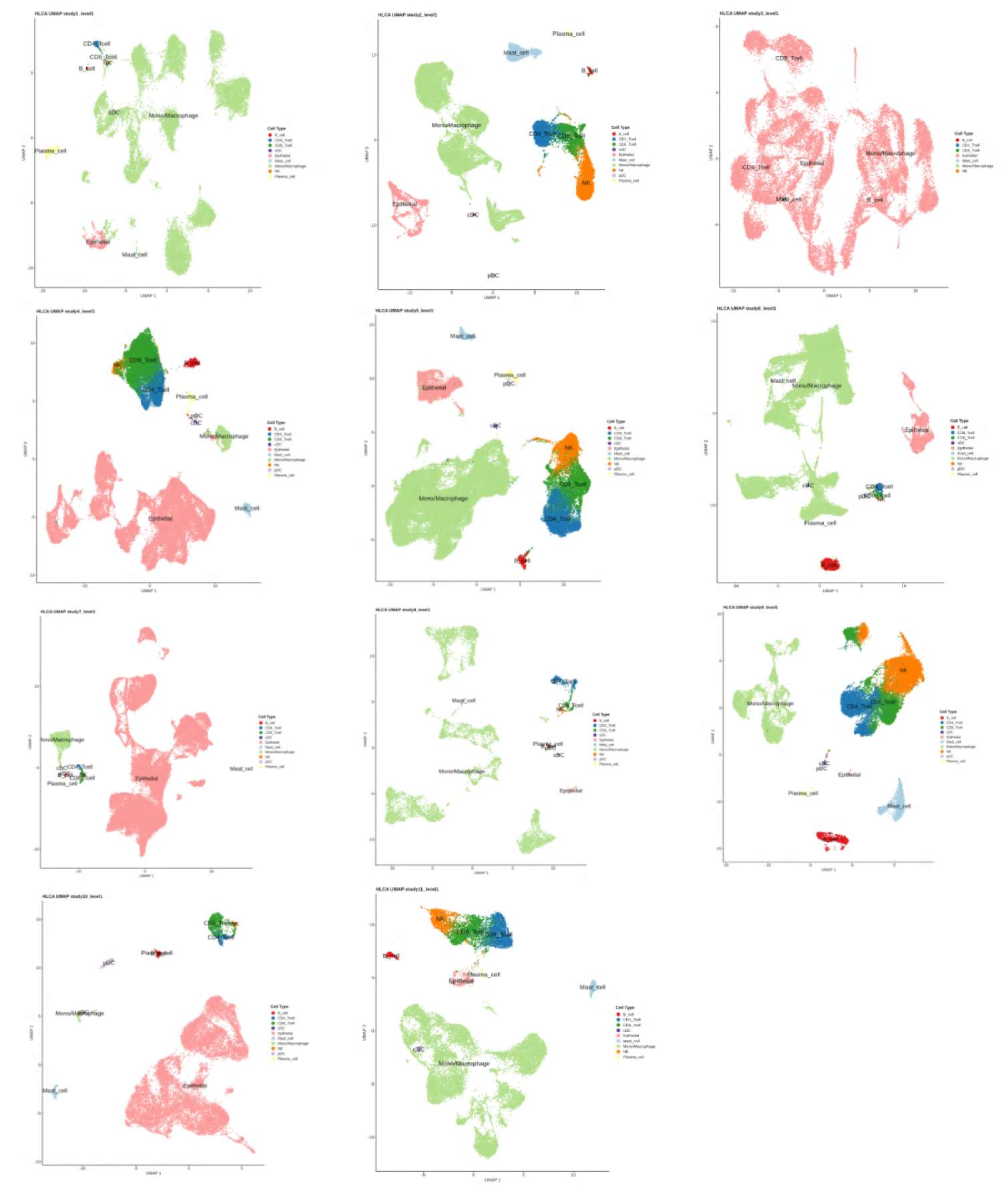
UMAPs of Level-1 cell type annotations across the 11 studies in the HLCA core dataset. Each panel shows a UMAP embeddings for one study included in the analysis. Colors denote annotated Level-1 cell types retrieved from the HLCA dataset, and labels indicate corresponding cell type names. Cell type proportions and distributions vary across these scRNA-seq studies within the HLCA core dataset.

**Fig. S2.**
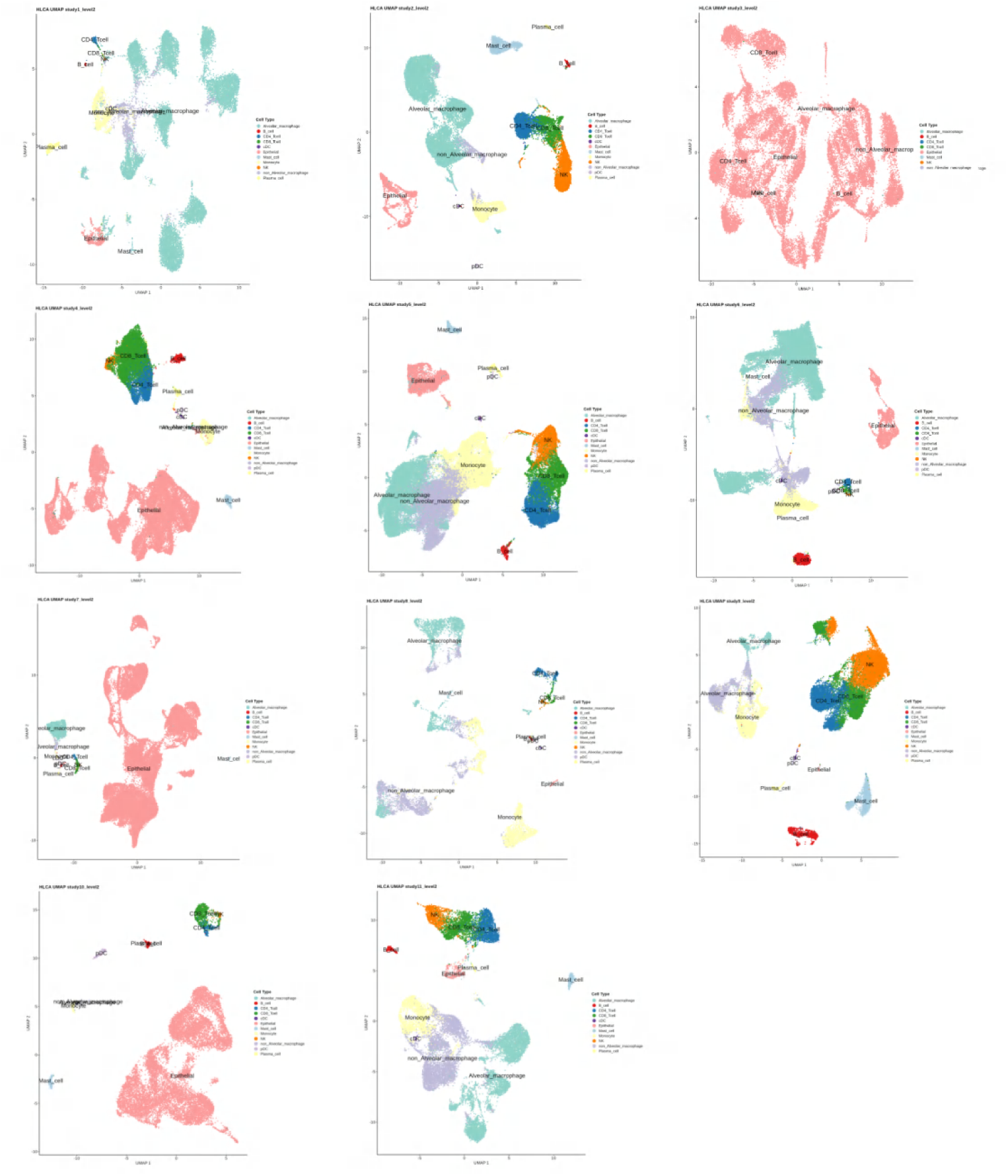
UMAPs of Level-2 cell type annotations across the 11 studies in the HLCA core dataset. Each panel shows a UMAP embeddings for one study included in the analysis. Colors denote annotated Level-2 cell types retrieved from the HLCA dataset, and labels indicate corresponding cell type names. Cell type proportions and distributions vary across these scRNA-seq studies within the HLCA core dataset.

**Fig. S3.**
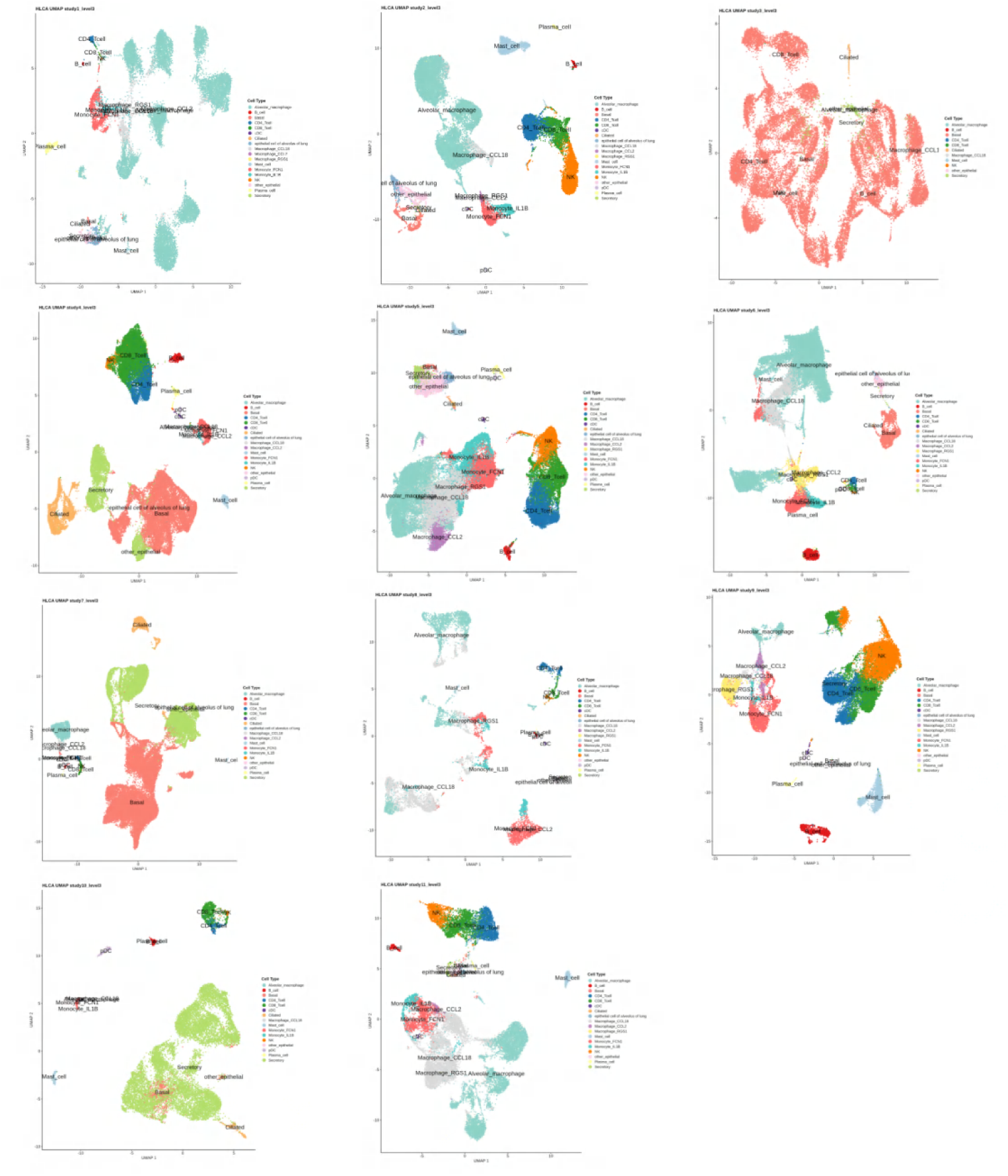
UMAPs of Level-3 cell type annotations across the 11 studies in the HLCA core dataset. Each panel shows a UMAP embeddings for one study included in the analysis. Colors denote annotated Level-3 cell types retrieved from the HLCA dataset, and labels indicate corresponding cell type names. Cell type proportions and distributions vary across these scRNA-seq studies within the HLCA core dataset.

**Fig. S4.**
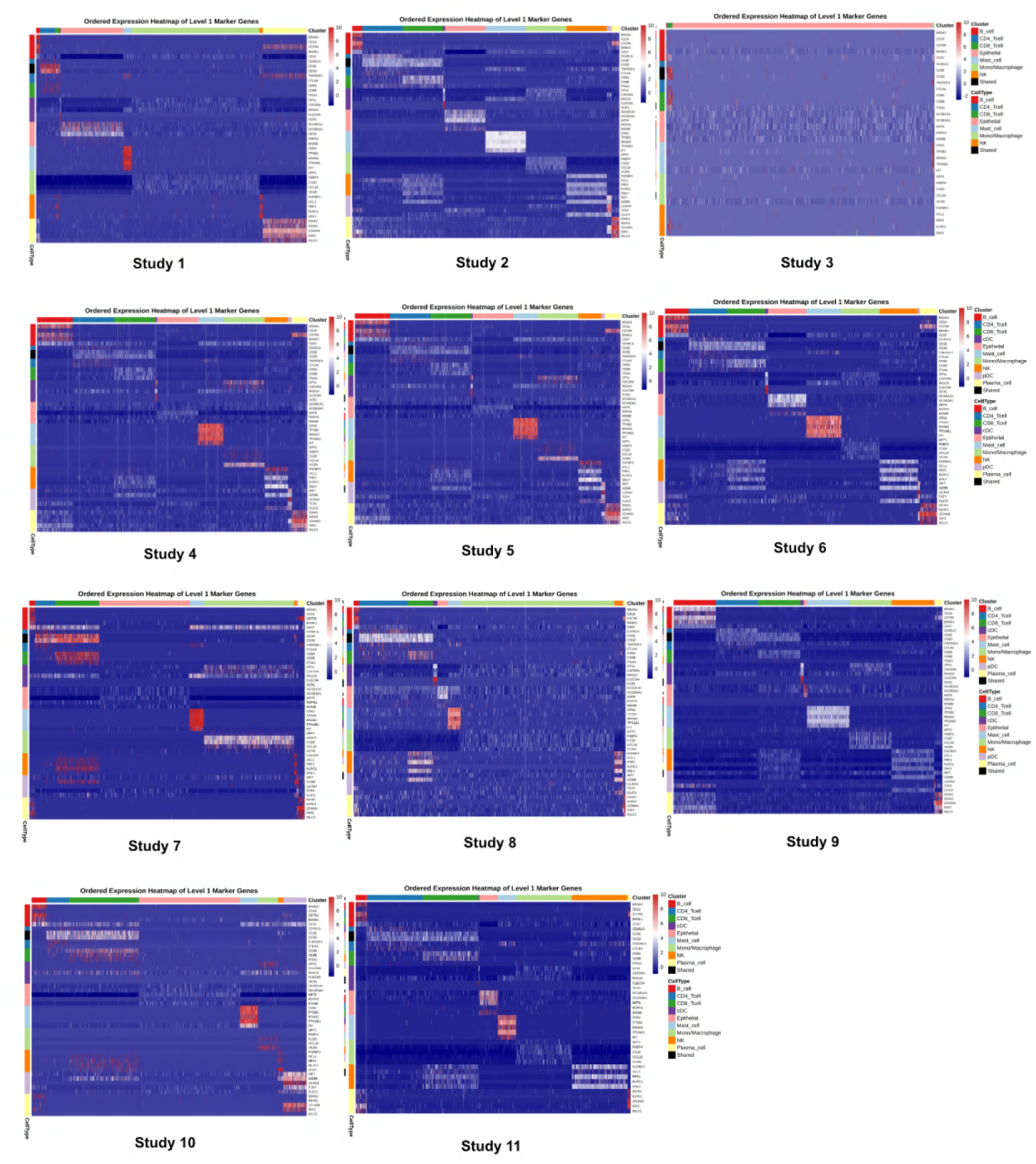
Heatmap of the expression of top Level-1 cell-type-specific marker genes across scRNA-seq studies in the HLCA core dataset.

**Fig. S5.**
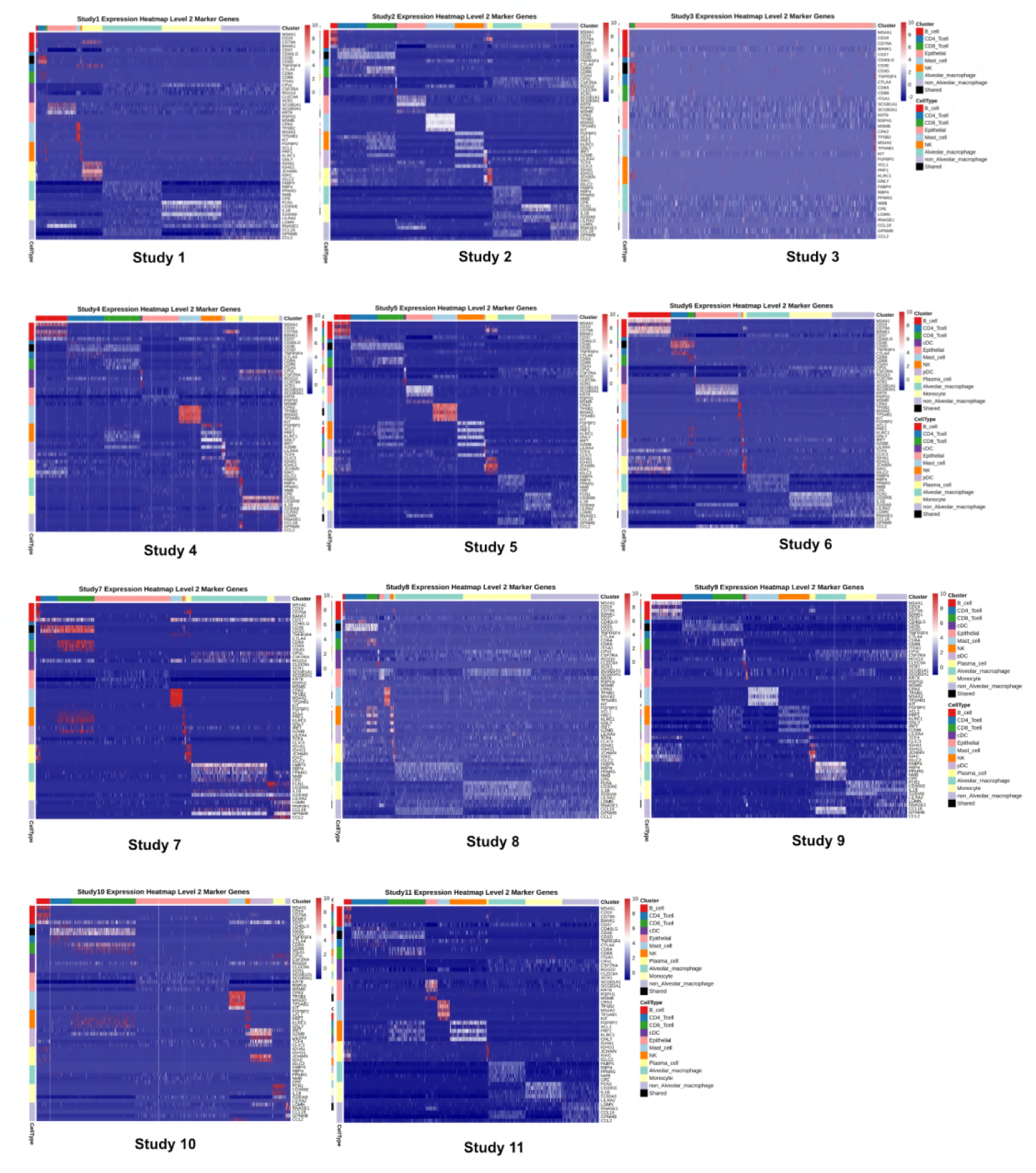
Heatmap of the expression of top Level-2 cell-type-specific marker genes across scRNA-seq studies in the HLCA core dataset.

**Fig. S6.**
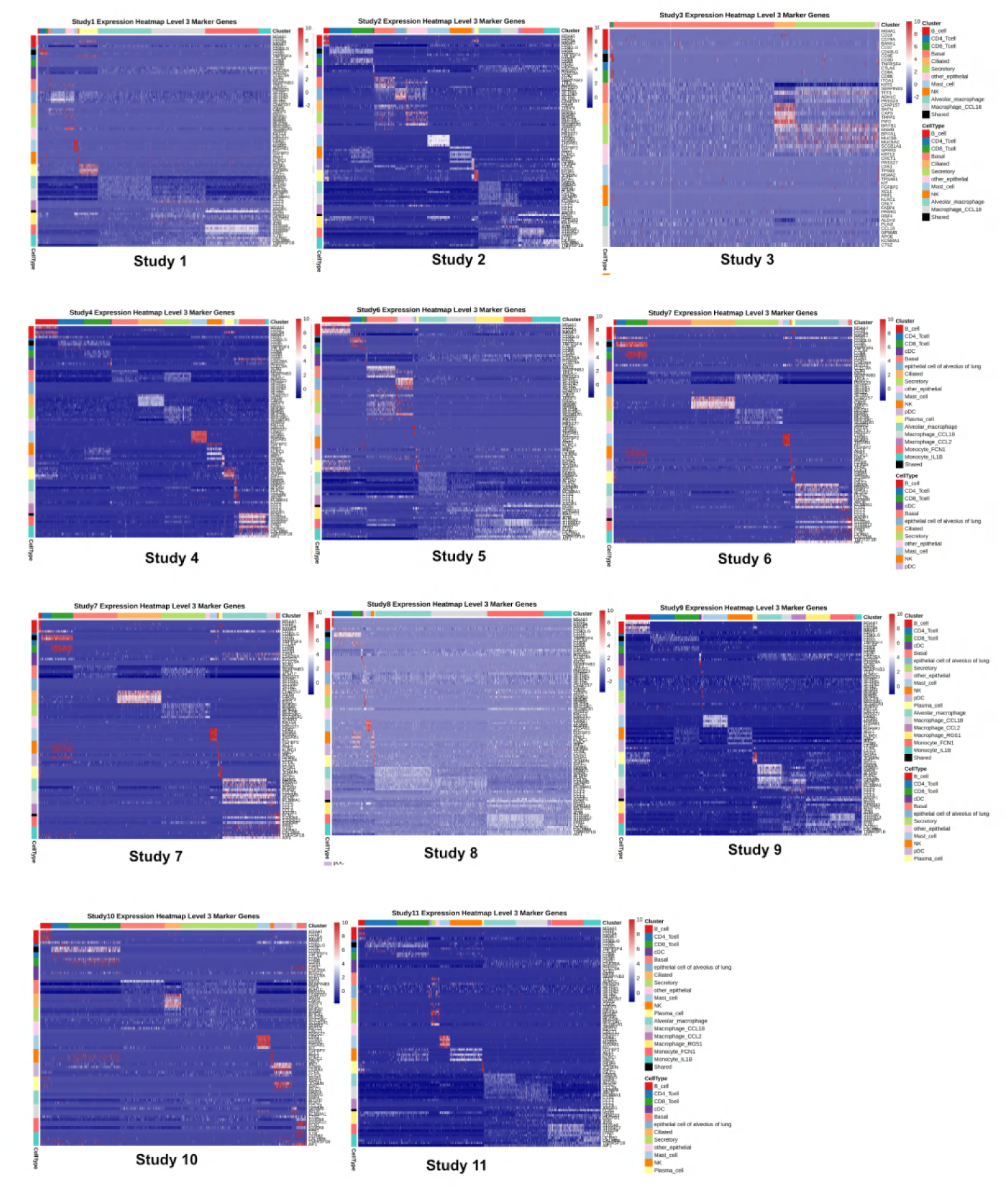
Heatmap of the expression of top Level-3 cell-type-specific marker genes across scRNA-seq studies in the HLCA core dataset.

**Fig. S7.**
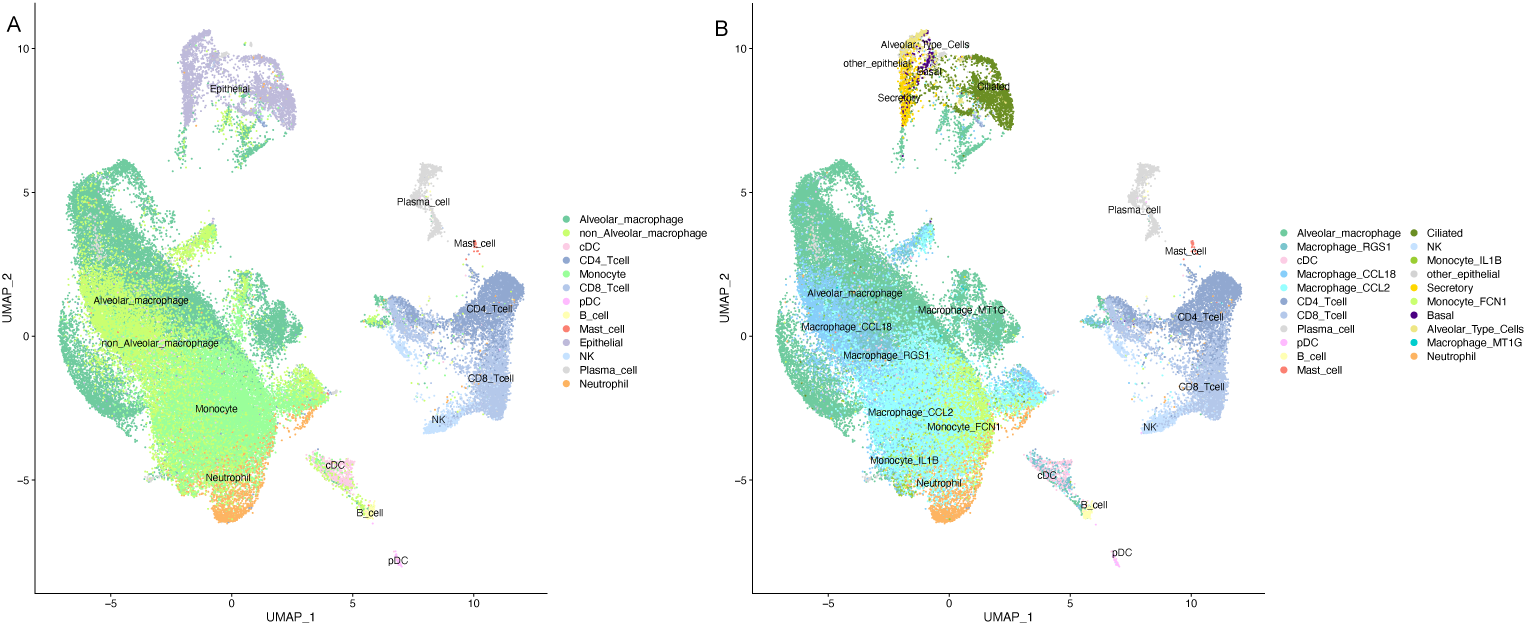
UMAPs plots of cell type annotation for the Liao *et. al* dataset with higher resolutions. (A). Cell type Level 2 annotations using BAL-EA. (B). Cell type Level 3 annotations using BAL-EA.

**Fig. S8.**
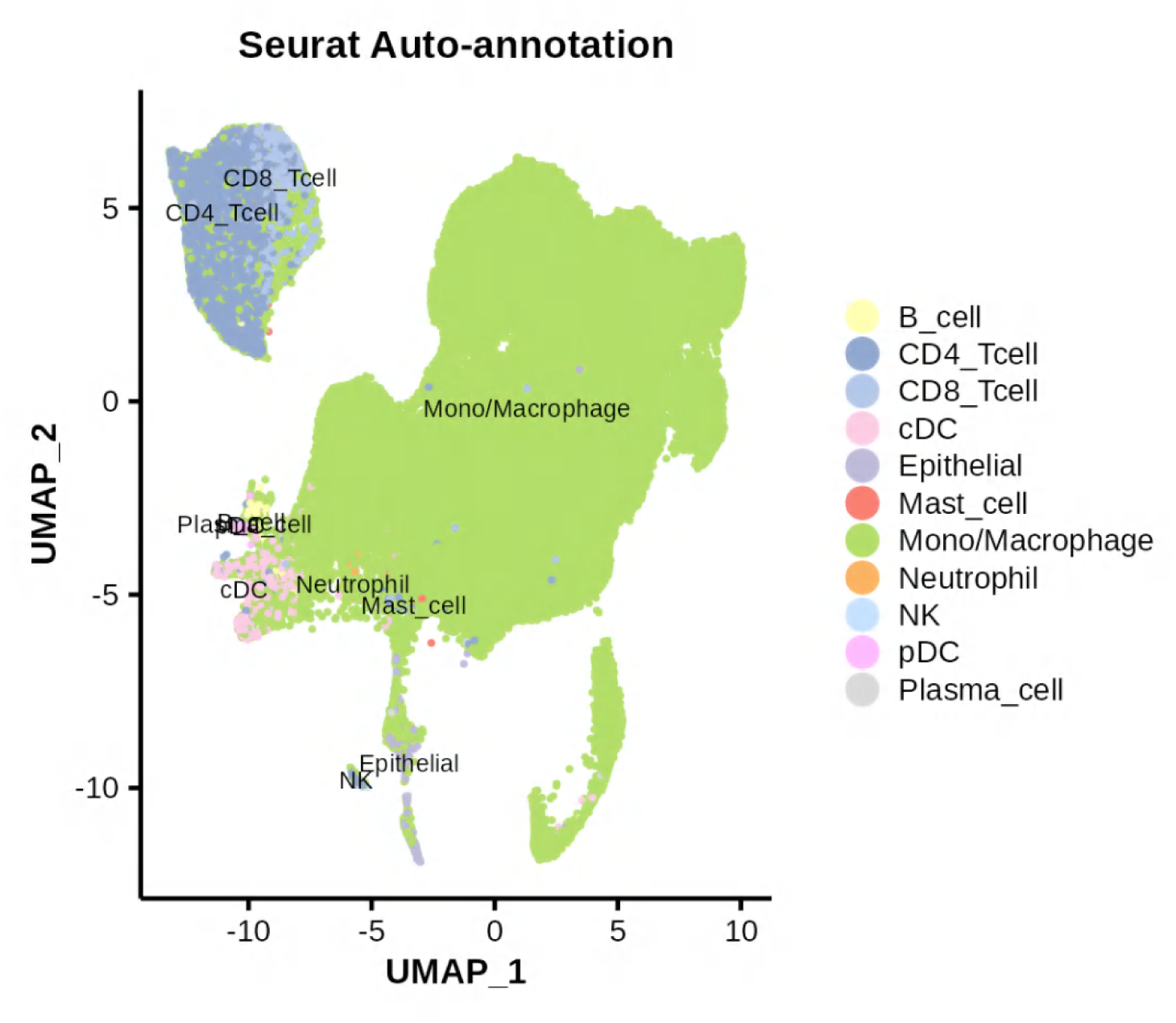
UMAPs plots of cell type annotation for the *in-house* dataset with Seurat cell type labels.

**Fig. S9.**
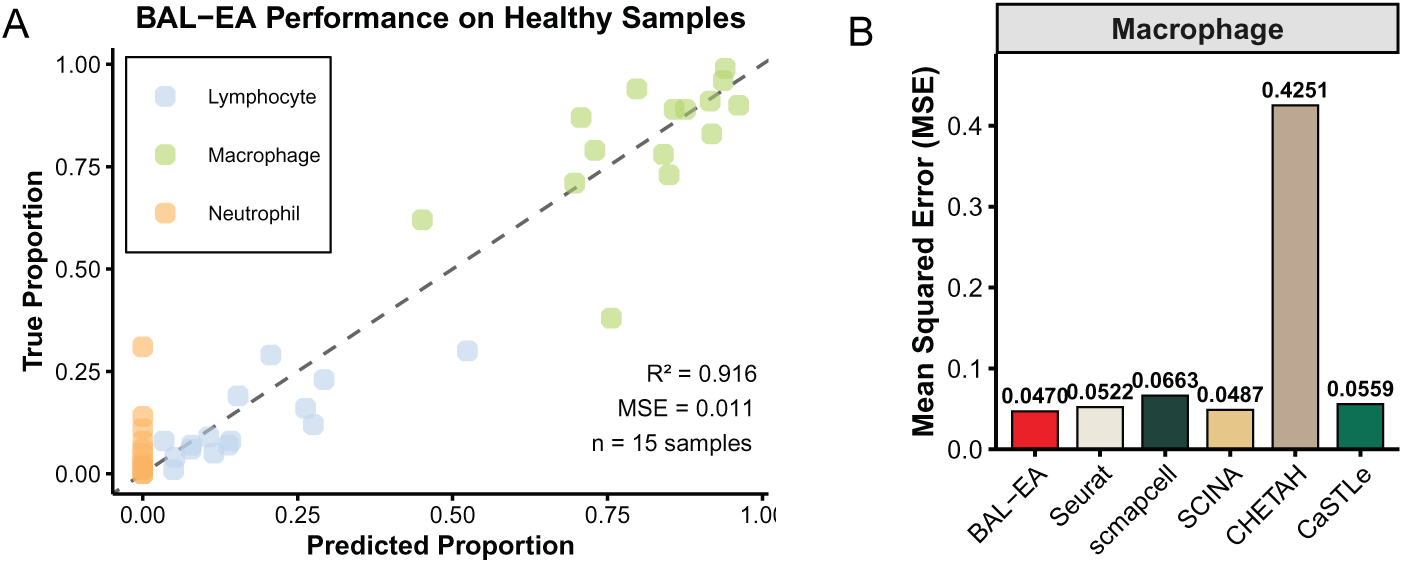
Performance validation of BAL-EA on healthy BAL samples. (A) Correlation between predicted and true cell type proportions across three major cell types in 15 healthy individuals (R2 = 0.916, MSE = 0.011). (B) Comparison of macrophage quantification accuracy across six autoannotation methods, showing BAL-EA achieves the lowest mean squared error (MSE = 0.047).

**Fig. S10.**
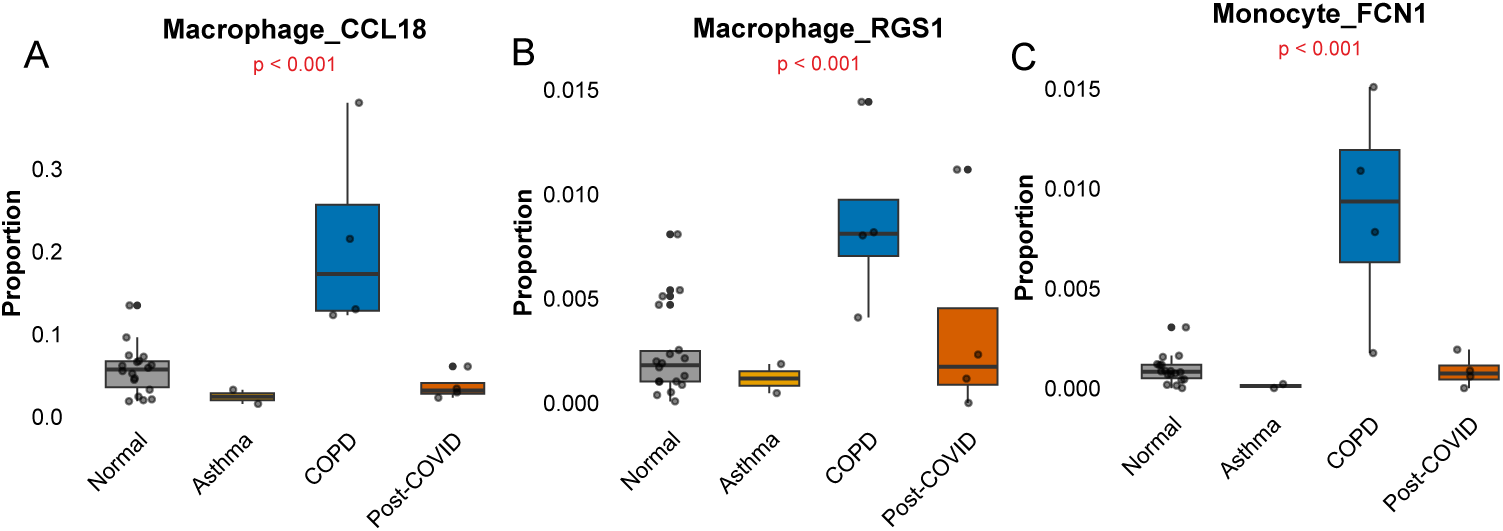
Disease-associated changes in proinflammatory macrophage and monocyte subsets across respiratory conditions. Boxplots showing proportions of (A) macrophages CCL18, (B) macrophages RGS1, and (C) monocytes FCN1. P-values from binomial GLM adjusted for age and sex; p *<* 0.001: COPD vs Normal.

## References

[1] Ernst, A. & Herth, F. J. F. (eds) Principles and Practice of Interventional Pulmonology (Springer, New York, NY, 2013).

[2] Gerayeli, F. V. et al. Single-cell RNA sequencing of bronchoscopy specimens: development of a rapid minimal handling protocol. BioTechniques 75, 157–167 (2023).

[3] Sin, D. D. What Single Cell RNA Sequencing Has Taught Us about Chronic Obstructive Pulmonary Disease. Tuberculosis and Respiratory Diseases 87, 252–260 (2024).

[4] Liao, M. et al. Single-cell landscape of bronchoalveolar immune cells in patients with COVID-19. Nature Medicine 26, 842–844 (2020).

[5] Wauters, E. et al. Discriminating mild from critical COVID-19 by innate and adaptive immune single-cell profiling of bronchoalveolar lavages. Cell Research 31 (2021).

[6] Grant, R. A. et al. Circuits between infected macrophages and T cells in SARS-CoV-2 pneumonia. Nature 590, 635–641 (2021).

[7] He, J. et al. Single-cell analysis reveals bronchoalveolar epithelial dysfunction in COVID-19 patients. Protein & Cell 11, 680–687 (2020).

[8] Chua, R. L. et al. COVID-19 severity correlates with airway epithelium-immune cell interactions identified by single-cell analysis. Nature Biotechnology 38, 970–979 (2020).

[9] Li, X. et al. ScRNA-seq expression of IFI27 and APOC2 identifies four alveolar macrophage superclusters in healthy BALF. Life Science Alliance 5 (2022).

[10] Mould, K. J. et al. Airspace Macrophages and Monocytes Exist in Transcriptionally Distinct Subsets in Healthy Adults. American Journal of Respiratory and Critical Care Medicine 203, 946–956 (2021).

[11] Hu, Y. et al. Single-cell sequencing of lung macrophages and monocytes reveals novel therapeutic targets in copd. Cells 12, 2771 (2023).

[12] Sikkema, L. et al. An integrated cell atlas of the lung in health and disease. Nature Medicine 29, 1563–1577 (2023).

[13] Aran, D. et al. Reference-based analysis of lung single-cell sequencing reveals a transitional profibrotic macrophage. Nature Immunology 20, 163–172 (2019).

[14] Kiselev, V. Y., Yiu, A. & Hemberg, M. scmap: projection of single-cell RNA-seq data across data sets. Nature Methods 15, 359–362 (2018).

[15] Lieberman, Y., Rokach, L. & Shay, T. CaSTLe – Classification of single cells by transfer learning: Harnessing the power of publicly available single cell RNA sequencing experiments to annotate new experiments. PLoS ONE 13, e0205499 (2018).

[16] de Kanter, J. K., Lijnzaad, P., Candelli, T., Margaritis, T. & Holstege, F. C. P. CHETAH: a selective, hierarchical cell type identification method for single-cell RNA sequencing. Nucleic Acids Research 47, e95 (2019).

[17] Zhang, Z. et al. SCINA: A Semi-Supervised Subtyping Algorithm of Single Cells and Bulk Samples. Genes 10, 531 (2019).

[18] Abdelaal, T. et al. A comparison of automatic cell identification methods for single-cell rna sequencing data. Genome Biology 20, 194 (2019).

[19] Morse, C. et al. Proliferating SPP1/MERTK-expressing macrophages in idiopathic pulmonary fibrosis. The European Respiratory Journal 54, 1802441 (2019).

[20] Bai, K., Moa, B., Shao, X. & Zhang, X. Pclda: An interpretable cell annotation tool for single-cell rna-sequencing data based on simple statistical methods. Computational and Structural Biotechnology Journal 27, 3264–3274 (2025).

[21] Hao, Y. et al. Integrated analysis of multimodal single-cell data. Cell 184, 3573–3587 (2021).

[22] Kiselev, V. Y., Andrews, T. S. & Hemberg, M. Challenges in unsupervised clustering of single-cell RNA-seq data. Nature Reviews Genetics 20, 273–282 (2019).

[23] Luecken, M. D. & Theis, F. J. Current best practices in single-cell RNA-seq analysis: a tutorial. Molecular Systems Biology 15, e8746 (2019).

[24] Xu, C. et al. Probabilistic harmonization and annotation of single-cell transcriptomics data with deep generative models. Molecular Systems Biology 17, e9620 (2021).

[25] Wolf, F. A., Angerer, P. & Theis, F. J. SCANPY: large-scale single-cell gene expression data analysis. Genome Biology 19, 15 (2018).

[26] Wolock, S. L., Lopez, R. & Klein, A. M. Scrublet: Computational Identification of Cell Doublets in Single-Cell Transcriptomic Data. Cell systems 8, 281–291.e9 (2019).

